# Identification and characterization of innate immunity in *Actinidia melanandra* in response to *Pseudomonas syringae* pv. *actinidiae*

**DOI:** 10.1101/2024.04.16.589833

**Authors:** Lauren M. Hemara, Abhishek Chatterjee, Shin-Mei Yeh, Ronan K.Y. Chen, Elena Hilario, Liam Le Lievre, Ross N. Crowhurst, Deborah Bohne, Saadiah Arshed, Haileigh R. Patterson, Kelvina Barrett-Manako, Susan Thomson, Andrew C. Allan, Cyril Brendolise, David Chagné, Matthew D. Templeton, Jay Jayaraman, Jibran Tahir

**Affiliations:** School of Biological Sciences, The University of Auckland, Auckland, New Zealand; The New Zealand Institute for Plant and Food Research Limited, Mount Albert Research Centre, New Zealand; The New Zealand Institute for Plant and Food Research Limited, Palmerston North, New Zealand; Department of Biochemistry, University of Otago, Dunedin, New Zealand; The New Zealand Institute for Plant and Food Research Limited, Lincoln Research Centre, New Zealand

## Abstract

*Pseudomonas syringae* pv. *actinidiae* biovar 3 (Psa3) has decimated kiwifruit orchards growing susceptible kiwifruit *Actinidia chinensis* varieties. Effector loss has occurred recently in Psa3 isolates from resistant kiwifruit germplasm, resulting in strains capable of partially overcoming resistance present in kiwiberry vines (*A. arguta, A. polygama*, and *A. melanandra*). Diploid male *A. melanandra* recognises several effectors, sharing recognition of at least one avirulence effector (HopAW1a) with previously studied tetraploid kiwiberry vines. Sequencing and assembly of the *A. melanandra* genome enabled the characterisation of the transcriptomic response of this non-host to wild-type and genetic mutants of Psa3. *A. melanandra* appears to mount a classic effector-triggered immunity (ETI) response to wildtype Psa3 V-13, as expected. Surprisingly, the type III secretion (T3S) system-lacking *Psa3 V-13 ΔhrcC* strain did not appear to trigger pattern-triggered immunity (PTI) despite lacking the ability to deliver immunity-suppressing effectors. Contrasting the *A. melanandra* responses to an effectorless Psa3 V-13 Δ*33E* strain and to Psa3 V-13 Δ*hrcC* suggested that PTI triggered by Psa3 V-13 was based on the recognition of the T3S itself. The characterisation of both ETI and PTI branches of innate immunity responses within *A. melanandra* further enables breeding for durable resistance in future kiwifruit cultivars.

## Introduction

Kiwifruit (*Actinidia* spp.) is a valuable perennial crop threatened by the bacterial pathogen *Pseudomonas syringae* pv. *actinidiae* (Psa) (McCann et al., 2017; Scortichini et al., 2023). Psa biovar 3 (Psa3) spread throughout kiwifruit-growing regions worldwide during a pandemic in the late 2000s, causing significant economic losses. In New Zealand, Psa3 V-13 (ICMP 18884) represents the initial Psa3 incursion that decimated orchards growing a monoculture of the highly susceptible cultivar *Actinidia chinensis* var. *chinensis* ‘Hort16A’. Replacing ‘Hort16A’ with less susceptible cultivars has helped the New Zealand kiwifruit industry recover from the impact of this disease since the initial incursion. However, Psa remains a persistent challenge, requiring significant time and expense to control through chemical applications and orchard hygiene practices. The use of copper-based sprays was initially effective in managing Psa. However, this has led to the wide-spread emergence of copper-tolerant Psa3 strains through the acquisition of copper resistance genes on integrative conjugative elements and plasmids, increasing Psa’s copper tolerance from an effective “naïve” minimum inhibitory concentration (MIC) of 0 mM CuSO_4_ to over 1.6 mM (Colombi et al., 2017). To sustainably manage Psa long-term, there is a need to diversify Psa management strategies and to develop durable Psa-resistant kiwifruit cultivars.

To breed durable pathogen resistance in crops, it is important to capture the causal genetic loci for diverse modes of plant immunity. Previously, it was shown within the commercially dominant *A. chinensis* species complex that Psa tolerance and susceptibility are mostly associated with multiple loci in yellow-fleshed tetraploid and diploid outcrosses in field conditions (Tahir et al., 2019, 2020). Some degree of Psa resistance has been observed within the *A. chinensis* species complex. Surveys suggest that other *Actinidia* species in the Leiocarpae section, including *A. macrosperma, A. valvata, A. polygama, A. hypoleuca, A. arguta*, and *A. melanandra* could potentially be Psa-resistant, with fewer vines removed from the Te Puke Research Orchard germplasm collection in New Zealand because of Psa infection (Datson et al., 2015). These potentially Psa-resistant species also fall clearly within the monophyletic smooth-skinned fruit (SSF) clade (Liu et al., 2017). *A. melanandra* is native to the Hubei and Yunnan provinces of China, has purple-red kiwiberry fruit, and is of particular interest owing to several diploid genotypes being identified (Datson et al., 2015). Psa3 symptoms have previously been observed on a commercial *A. arguta* orchard; however, both the appearance of symptoms and the impact on orchard production was very limited (Vanneste et al., 2014). Inter-specific hybridisation in kiwifruit will open doors to more robust genetic combinations against the more rapidly evolving pathogen (Z. Wang et al., 2017).

Plant innate immunity is a complex process that consists of two interconnected layers, namely pattern-triggered immunity (PTI) and effector-triggered immunity (ETI). PTI is triggered by the recognition of highly conserved pathogen-associated molecular patterns at the host cell membrane. ETI is a more specific response that is activated by the presence or activity of one or more pathogen effector proteins, typically within the host cell. ETI potentiates and amplifies PTI, often culminating in the hypersensitive response, leading to the death of infected cells to protect the rest of the plant (Ngou et al., 2021; Yuan et al., 2021). While ETI can be highly effective in protecting plants from disease, resistance mediated by a single resistance (R) gene may be vulnerable to resistance breakdown because of pathogen evolution. R gene stacking may increase the durability of resistance in the field by providing multiple avenues of pathogen recognition that are harder for the pathogen to evade.

In Psa-resistant kiwiberry *A. arguta* versus Psa-susceptible *A. chinensis*, ETI and PTI responses can be affected by the presence or absence of avirulence and virulence Psa effectors (Hemara et al., 2022; Jayaraman et al., 2021, 2023). The molecular signatures underling these dynamically interconnected modes of host immunity can help identify pathways that are specifically activated and co-evolved in both compatible and incompatible kiwifruit-Psa interactions. In this current study, the emergence of Psa3 effector-variant strains on various host kiwifruit species was explored and a diploid kiwiberry model system (*A. melanandra*) was utilised to dissect layers of innate immune response to Psa3. By exploring the transcriptional landscape to infection by virulent and non-virulent strains of Psa3 and *P. fluorescens* (Pfo) in the diploid Psa3-resistant *A. melanandra*, ETI and PTI pathways in *Actinidia* were characterised. This will allow for the mapping of novel genes for resistance and breeding cultivars that will have durable, long-term resistance to Psa in the orchard.

## Results

### Psa3 lineages with deleted effectors have emerged independently and multiple times on Psa-resistant *Actinidia* vines

*hopAW1a* loss has been previously reported on *A. arguta*, as typified by Psa3 X-27 (Hemara et al., 2022). Following Psa3 genome bio-surveillance in the *Actinidia* germplasm collection at Te Puke, New Zealand from 2017 to 2022, the further independent emergence of two additional lineages with deletions in the exchangeable effector locus was observed (EEL; Fig. 1A). Most EEL-loss isolates have a 51 kb deletion mediated by recombination at DD[E/D] transposases (DDEs) and Miniature inverted-repeat transposable elements (MITEs). However, Psa3 X-469 has a smaller 38 kb deletion, which removes *hopQ1a, hopD1a, avrD1, avrB2b, hopAB1ak*, and *hopF4a* alongside *hopAW1a, hopF1e, hopAF1b, hopD2a*, and *hopF1a* (Fig. 1A; Fig. S1A). Unlike Psa3 X-27, Psa3 X-469 retains the NRPS toxin biosynthesis cluster. This deletion appears to be mediated by recombination at the flanking MITEg3 and MITEg8 elements (Fig. S1A). The emergence of two independent *avrRpm1a* loss lineages was also observed (Fig. 1A). The *avrRpm1a* deletion of 11 kb is flanked by Tn family 21 and DDEg14 (Fig. S1B). Both *hopAW1a* and *avrRpm1a* are known to be recognised by Psa-resistant *A. arguta* (Hemara et al., 2022). Interestingly, the loss of *hopZ5a* or *hopF1c* from isolates collected from these diverse germplasm vines was never observed. This is despite the fact that *hopZ5a* (like *avrRpm1a* that is lost from the orchard isolates) is recognized by *A. arguta* and is ‘cargo’ on an exapt ICE-A, a remnant, non-mobile integrative conjugative element that lacks a t-RNA-Lys attachment (att) site for excision (Hemara et al., 2022; Poulter et al., 2018). While the majority of these strains have been isolated from *A. arguta*, EEL loss isolates have also been found on *A. hypoleuca* and *A. melanandra* vines, and *avrRpm1a* loss isolates have been found on *A. polygama* and *A. glaucophylla* (Fig. 1A). *A. melanandra* is a species of interest because diploid members of this species possess multiple resistance loci against Psa3, and its simple ploidy makes it amenable to genome sequencing and a less complex genome assembly, and utility in mapping target loci for breeding.

**Figure 1.**
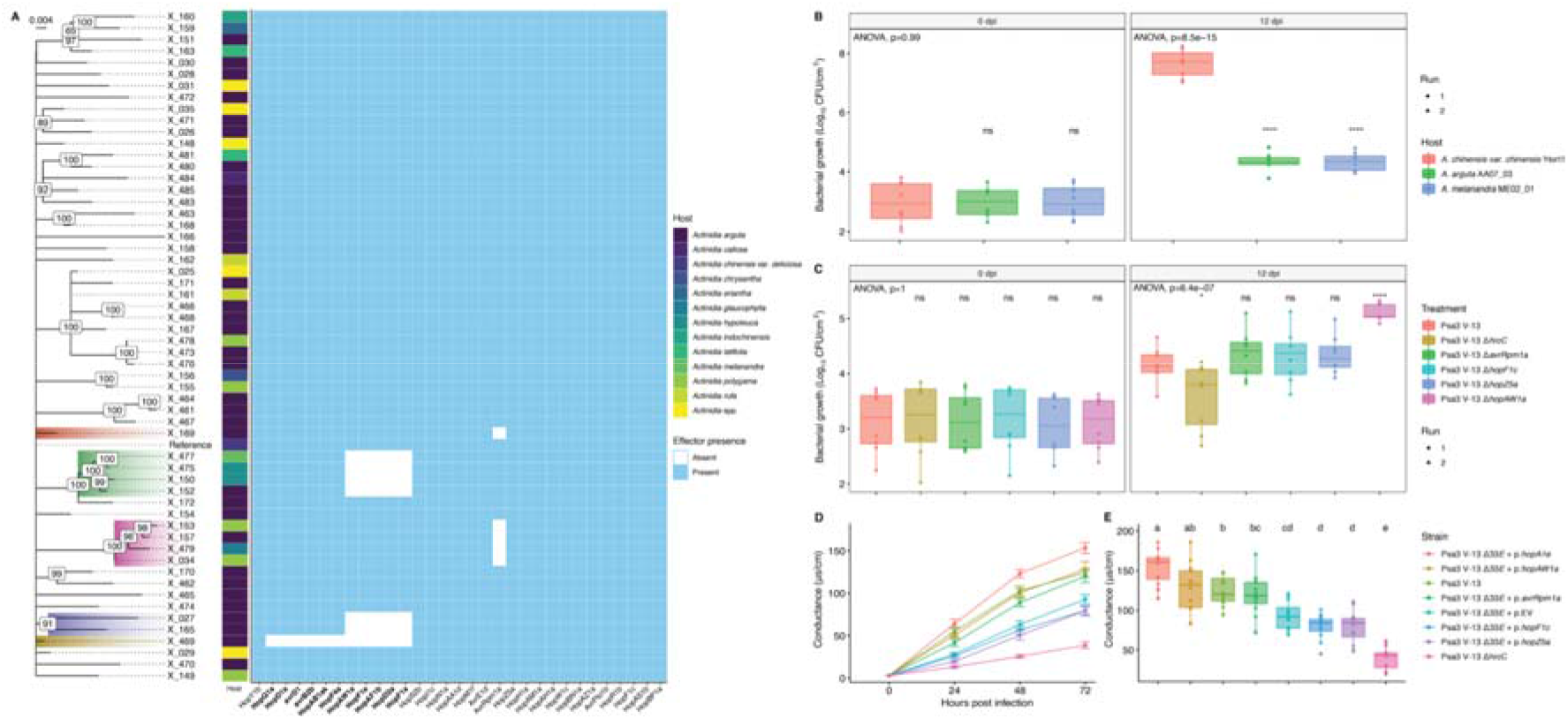
*Actinidia melanandra* mounts an effector-triggered defence response against *Pseudomonas syringae* pv. actinidiae (Psa3) V-13. (A) Core SNP phylogeny of Psa isolates from germplasm *Actinidia* vines. Effector presence and absence by genome position indicates the emergence of multiple lineages of exchangeable effector locus (EEL) and avrRpm1a loss variants. Effectors in the exchangeable effector locus are highlighted in bold. (B) *A. chinensis* var. *chinensis* ‘Hort16A’, *A. arguta* AA07_03, and *A. melanandra* ME02_01 plantlets were flood-inoculated with Psa3 V-13 at approximately 10^6^ CFU/mL. Bacterial growth was quantified relative to ‘Hort16A’ at 0-and 12-days post-inoculation by plate count. Box and whisker plots, with black bars representing the median values for the four pseudobiological replicates and whiskers representing the 1.5 inter-quartile range. Asterisks indicate the statistically significant difference of Student’s t-test between the indicated species and *A. chinensis* var. *chinensis* ‘Hort16A’, where *p*≤0.05 (*), *p*≤0.01 (**), *p*≤0.001 (***), and *p*>0.05 (ns; not significant). (C) *A. melanandra* ME02_01 plantlets were flood-inoculated with Psa3 V-13 effector knockout strains at approximately 10 CFU/mL. Bacterial growth was quantified relative to Psa3 V-13 at 0- and 12-days post-inoculation by plate count. Box and whisker plots, with black bars representing the median values for the four pseudobiological replicates and whiskers representing the 1.5 inter-quartile range. Asterisks indicate the statistically significant difference of Student’s t-test between the indicated strain and wild-type Psa3 V-13, where *p*≤0.05 (*), *p*≤0.01 (**), *p*≤0.001 (***), and *p*>0.05 (ns; not significant). (D-E) Leaf discs from *A. melanandra* ME02_01 plantlets were vacuum-infiltrated with Psa3 V-13 or Psa3 V-13 Δ33E carrying empty vector (EV) or a plasmid-borne type III secreted effector (*hopAW1a, hopZ5a*, avrRpm1a or *hopF1c*, or positive control *hopA1j* from P. *syringae* pv. *syringae* 61) inoculum at ∼5 x 10^8^ CFU/mL. (D) Electrical conductivity due to HR-associated ion leakage was measured at indicated times over 72 hours. The ion leakage curves are stacked for three independent runs of this experiment. Error bars represent the standard errors of the means calculated from the five pseudobiological replicates per experiment (n = 15). (E) Tukey’s HSD with different letters indicates treatment groups that are significantly different at the 72-h timepoint (α ≤ 0.05).

### *A. melanandra* recognises HopAW1a, HopBP1a, and AvrRpm1a

*A. melanandra* (accession ME02_01) restricts the bacterial growth of wild-type Psa3 V-13, similarly to *A. arguta*, suggesting that *A. melanandra* might also recognise Psa3 through ETI (Fig. 1B). DAB staining indicates the production of reactive oxygen species (ROS) in the presence of Psa3 V-13, but not Psa3 V-13 Δ*hrcC*, supporting the existence of ETI in this accession (Fig. S2). However, *A. melanandra* may not necessarily recognise the same effector profile that *A. arguta* does. When comparing knockout strains for the four effectors recognized by *A. arguta*, only Psa3 Δ*hopAW1a* grew better than wild-type Psa3 on *A. melanandra*, while Δ*avrRpm1a*, Δ*hopF1c*, and Δ*hopZ5a* strains were not significantly different from wild-type Psa3 (Fig. 1C). To observe ETI-associated ion leakage, *P. fluorescens* carrying a T3S from *P. syringae* pv. *syringae* 61 (Pfo(T3S); Thomas et al., 2009), was used to deliver individual Psa3 V-13 effectors into *A. melanandra* (Fig. S3). Only HopAW1a and HopBP1a (both weakly) triggered ion leakage in ME02_01, suggesting some effector recognition (Fig. S3). This result was confirmed by a reporter eclipse assay (Jayaraman et al., 2021), demonstrating that both HopAW1a and HopBP1a, like the avirulence effector control HopA1j from *P. syringae* pv *syringae* 61, are recognized in *A. melanandra* (Fig. S4).

Pfo(T3S) may not be able to express and deliver effectors from Psa in the full context of a suite of other potential pathogenicity factors. To mimic this more complete context and deliver individual effectors from Psa3, a complete effector knockout strain (Psa3 V-13 Δ*33E*) was generated that lacked all 33 expressed effectors. To confirm that the AA07_03-recognized avirulence effectors were not affected by their level of expression under the synthetic promoter or the C-terminal HA tag, *hopAW1a, hopZ5a, avrRpm1a*, and *hopF1c* (with *shcF* carrying a point mutation resulting in an early truncation) were cloned under their native promoters. Psa3 V-13 Δ33E delivery of HopAW1a, HopZ5a, AvrRpm1a, and HopF1c revealed that HopAW1a and AvrRpm1a were able to trigger ion leakage in ME02_01 leaves, confirming resistance in *A. melanandra* ME02_01 is due to ETI (Fig. 1D–E). Curiously, Psa3 Δ*33E* + EV also appeared to trigger some ion leakage, despite lacking any functional effectors from Psa3 (Fig. 1D–E). This was not the case for Psa3 V-13 Δ*hrcC*, which, as expected, did not trigger ion leakage owing to its inability to secrete effectors in the absence of the type III secretion system (Fig. 1D–E). Pfo(T3S) + EV also caused some ion leakage in ME02_01, similarly to Psa3 Δ*33E*, and interestingly this response was also found to be completely abolished by Pfo(T3S) strains carrying some Psa effectors like *hopF1c, hopH1a* and *hopAZ1a* (Fig. S3).

### *A. melanandra* genome structure is orthologous to *A. arguta*

The diploid and Psa-resistant *A. melanandra* (ME02_01 accession) is a more suitable model system than the previously studied tetraploid *A. arguta* AA07_03 (Hemara et al., 2022) for both downstream utility in breeding and functional characterisation of trait genetics. Using Illumina short-read and Oxford Nanopore PromethION long-read platforms, the genome of *A. melanandra* (accession ME02_01) was sequenced and assembled. The genome contained 654.4 Mb from 728 scaffolds with an N50 of ∼ 21.2 Mb (comprising 1,746 contigs with N50 ∼1.4 Mb), and a Benchmarking Universal Single-Copy Orthologs (BUSCO) score of 98.9%. This assembly is referred to as ME02_01_v2.1, (WGS accession: JBAMMV000000000) from which 637.8 Mb was assigned to 29 chromosomes, aligning Hi-C reads. The gene models from the assembly were used to generate a draft gene set of 37,047 genes, of which 36,326 (98%) were present on the chromosomes. The final assembly and basic metrics of units (primary, unassigned, gene contents) are provided in Supplementary Data S1. Phylogenetic analyses of the *Actinidia* species have already shown that *A. arguta* and *A. melanandra* are the closest species among all taxa in the *Actinidia* clade (Tahir et al., 2022). The comparative whole-genome analysis between the publicly available diploid *A. arguta* var. *hypoleuca* (Akagi et al., 2023) and *A. melanandra* genomes show a high conservation of synteny between both species (Fig. 2). Notably, the male-specific Y-linked region in *A. melanandra* is also found on chr3 (11.32-11.63 Mb) as in *A. arguta* var. *hypoleuca*, instead of chr25 as in cultivated *A. chinensis* species (Akagi et al., 2023). This is the first comprehensive chromosome-scale genome representing the *A. melanandra* species.

**Figure 2.**
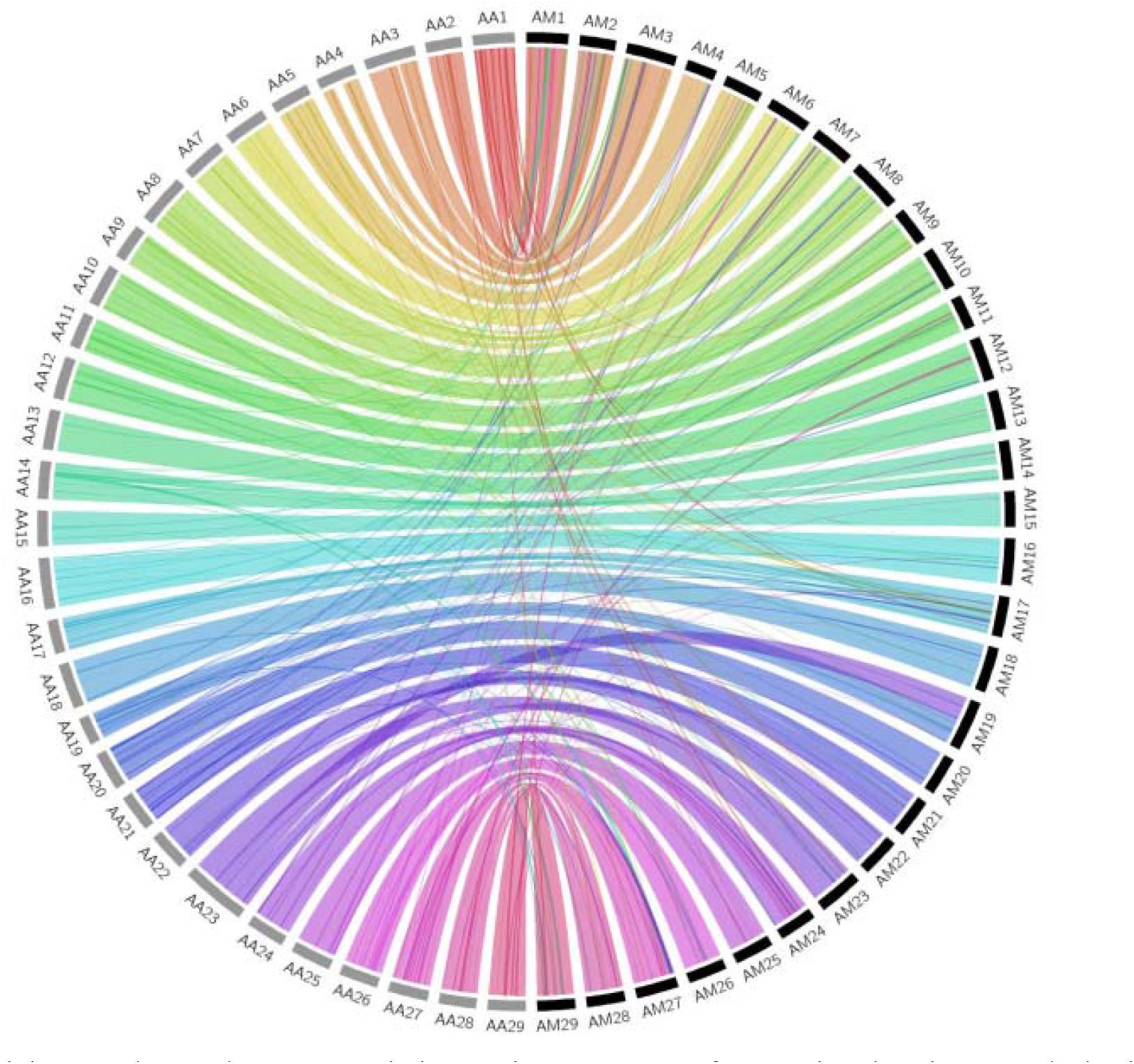
Whole genome (DNA:DNA) synteny Circos plot between chromosomes 1 to 29 from *Actinidia melanandra* ME02_01 (AM) and diploid *A. arguta var. hypoleuca* (AA) genome (Akagi et al., 2023). All by all alignments were performed using nucmer as described in the Methods and dnadiff used to filter ‘1-to-1 coordinates’. Coordinates were converted to Circos links using bundlelinks (-max_gap 1000000 - min_bundle_size 10000). Link bundles were coloured according to chromosome of origin.

### The ETI response of *A. melanandra* to Psa3

To dissect pathways of PTI and ETI in *A. melanandra*, a time-course experiment was performed where axenically grown ME02_01 plants were infiltrated with Psa3 V-13, Psa3 V-13 Δ*hrcC*, or buffer alone (mock). Leaf samples were harvested at designated timepoints for RNA extraction and expression analyses (Fig. S5). Gene expression (CPM) from all samples and treatments showed no significant differences in counts or quality (Fig. S6). Across the full time-series of 0, 3, 6, 10, 20, 30, and 40 h-post infiltration (hpi), a total of 3576 Differentially expressed genes (DEGs) were identified between the Psa3 V-13 treatment versus mock (adjusted *p*-value <0.001, |log_2_ fold-change|>2). Principal components analysis and a heatmap visualisation of gene expression suggest that there were two subsets of responses: one triggered by Psa3 V-13 and the other by mock and Psa3 V-13 Δ*hrcC* (Fig. 3A and B). The early timepoints (3 to 6 hpi) are crucial for cataloguing PTI responses (expected for both Psa3 V-13 and Psa3 V-13 Δ*hrcC*, but not mock) in common with and diverging from the ETI pathway. Meanwhile the mid timepoints (10 to 20 hpi) reflect the threshold time expected for ETI-specific expression, with an ETI response expected for Psa3 V-13 only (Fig. S5). Gene expression across the three treatments was clustered into five expression groups. Surprisingly, no gene clusters shared expression between Psa3 V-13 and Psa3 V-13 Δ*hrcC*, even though a strong gene expression response to Psa3 V-13 was observed within four out of five expression clusters, and only cluster 5 was strongly induced within the Psa3 V-13 Δ*hrcC* treatment (Fig. 3A–C).

**Figure 3.**
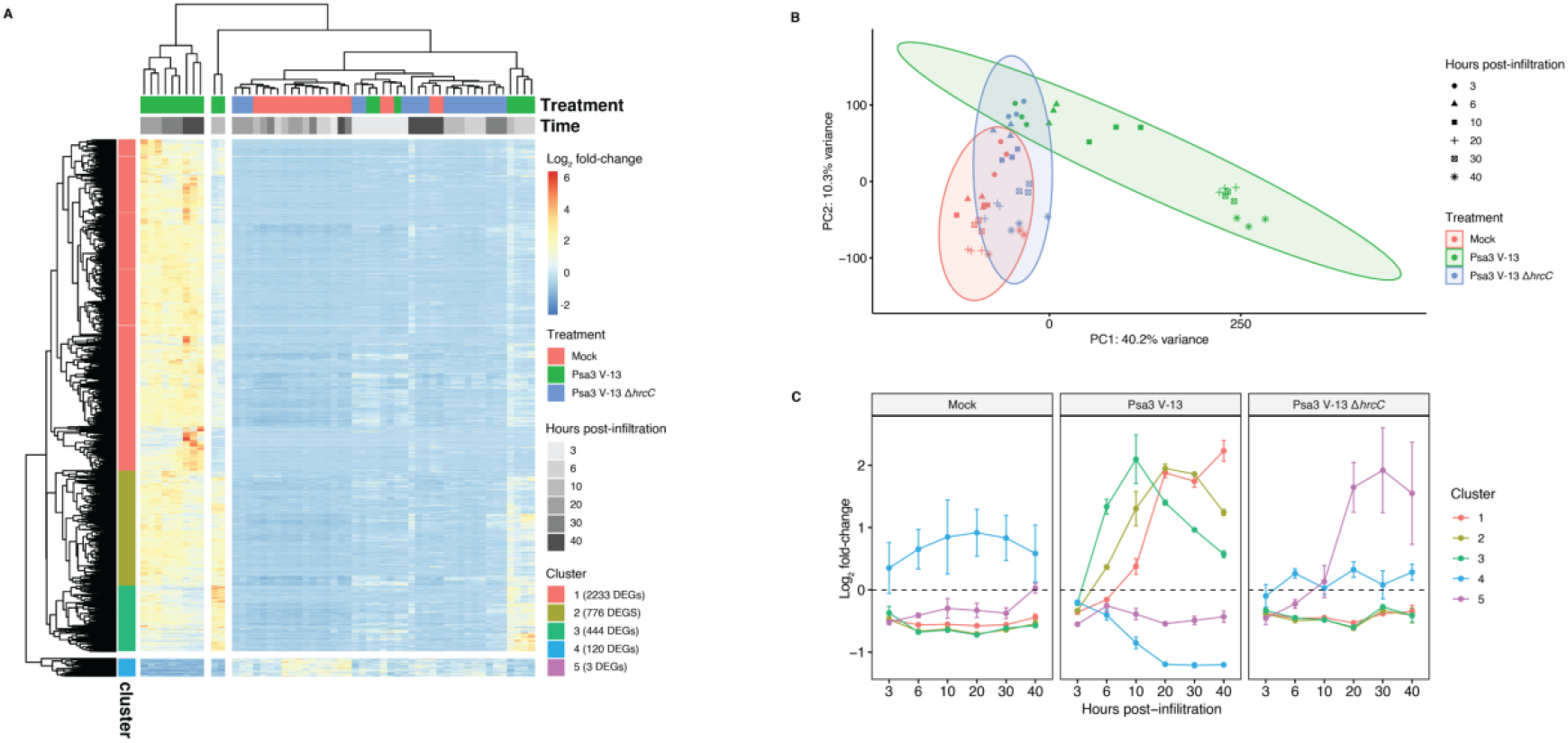
The transcriptional response to *Pseudomonas syringae pv. actinidiae* (Psa3) V-13 treatment in *Actinidia melanandra* ME02_01 over time. (A) Heatmap of differentially expressed genes induced by Psa3 V-13 treatment relative to mock (adjusted *p*-value <0.001). For each gene, raw counts were transformed by median of ratios normalization and Z-score scaling. The pheatmap package was used to generate a heatmap, with genes divided into five clusters based on gene expression patterns using the hclust() function. (B) Principal components analysis of gene expression as log_2_ fold change for all samples under all treatments and timepoints. (C) Mean log_2_ fold-change for each gene expression cluster over the 40-hour time series. Error bars indicate standard errors.

For Psa3 V-13 treatment, DEGs in cluster 3 (early Psa3-induced response) peaked earliest at 10 hpi, whereas cluster 2 DEGs (mid Psa3-induced response) peaked at 20 hpi, with cluster 1 DEGs (late Psa3-induced response) continuing to increase even at 40 hpi. Cluster 4 (Psa3-suppressed response) and cluster 5 (Psa3 Δ*hrcC*-induced response) represent a significantly smaller number of DEGs. Unsurprisingly, different subsets of defence-associated phosphorylation-regulating serine-threonine protein kinases were upregulated throughout the responses to Psa3 (Fig. 4A–C). Key defence mediator genes including WRKY transcription factors, mitogen activated protein kinases (MAPKs), ABC transporters, and cytochrome p450s were abundantly expressed at late timepoints (Fig. 4A). Gene families encoding putative E3 ubiquitin ligases, calcium binding proteins, and secretion-associated exocyst components were among the genes upregulated at early to mid-timepoints in response to Psa3 (Fig. 4B and C). Various classic defence-response marker genes including phenylalanine ammonia-lyases (PALs), WRKY and ERF transcription factors, pathogenesis-related proteins, flavin-dependent monooxygenases, and receptor kinase coding genes were upregulated at early to late timepoints specifically in response to Psa3 V-13 (Fig. S7). Interestingly, one of the DEGs with the highest level of expression was in cluster 1 (Fig. 4A) and encodes a GRIM-REAPER-like protein which, in *Arabidopsis*, binds to a receptor kinase to trigger ROS-induced cell death (Wrzaczek et al., 2009, 2015). Meanwhile, Psa3 V-13 triggered the suppression of a subset of genes including RNA recognition and stomatal closure-related proteins (Fig. 4D).

**Figure 4.**
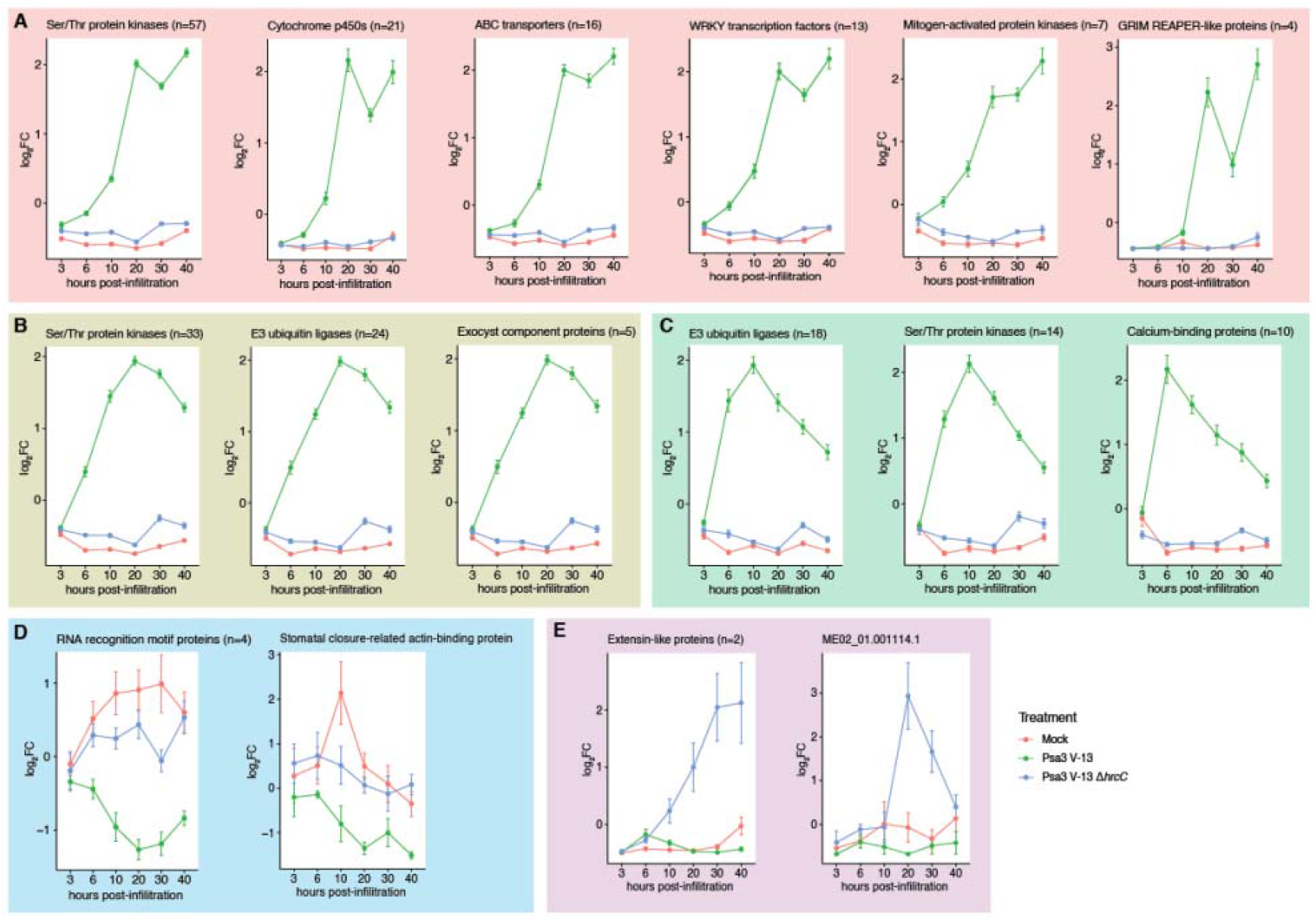
Expression of key families of differentially expressed genes in *Actinidia melanandra* ME02_01 for each treatment and timepoint in (A) early *Pseudomonas syringae* pv. *actinidiae* (Psa3)-induced response cluster 3, (B) mid Psa3-induced response cluster 2, (C) late Psa3-induced response cluster 1, (D) Psa3-suppressed response cluster 4, or (E) Psa3 Δ*hrcC* -induced response cluster 5. Expression for each family is averaged across the number of family members indicated and is plotted as log_2_ fold change for 3 replicates per sample across indicated timepoints (gene expression of family members passing the significance threshold *p*-value <0.001).

Surprisingly, Psa3 V-13 Δ*hrcC* triggered only a weak response using stringent criteria (adjusted *p*-value <0.001, |log_2_ fold-change|>2) with up-regulation of just three genes in cluster 5, including two defence-associated extension-like proteins (Fig. 4E). Excluding DEGs that respond to Psa V-13 and lowering the threshold (adjusted *p*-value <0.05) identified only twelve DEGs (with inconsistent expression between replicates) compared with the mock treatment, suggesting that *A. melanandra* is indeed unable to mount a significant defence response to Psa3 V-13 Δ*hrcC* (Fig. S8A and B).

### Psa3 Δ*33E* and Pfo(T3S) + *hopA1j* dependent PTI and ETI responses in *A. melanandra*

To further characterise *bona fide* ETI and PTI responses in *A. melanandra*, additional bacterial treatments were used in the experimental regime (Fig. S5). Sampling for these focused on early (3 and 6 hpi; PTI) and mid (20 hpi; ETI) timepoints, for examining gene expression responses. Gene expression (CPM) from these samples and treatments also showed no significant differences in counts or quality (Fig. S9). The three additional treatments included were: Psa3 Δ*33E* (lacking all predicted effectors), Pfo(T3S) + *hopA1j* (showing ETI in ME02_01, Fig. S2), and the non-virulent strain Pfo(T3S) + EV, and these were selected for comparison to the previous Psa3 and mock treatments. DEGs that are up-or downregulated (adjusted *p*-value <0.001, |log_2_ fold-change|>2) in each bacterial treatment, were filtered out across the three time points, from which genes were stacked across timepoints to display the cumulative response profile (Fig. 5A–E). Data from Psa3 V-13 and Psa3 V-13 Δ*hrcC* for the same timepoints were included, where ME02_01 showed a strong transcriptional response to wild-type Psa3 (with 2263 upregulated DEGs and 675 downregulated DEGs) and a significantly mild response to Psa3 V-13 Δ*hrcC* (with 169 upregulated DEGs and 50 downregulated DEGs) (Fig. 5A and B). The strong ion-leakage in ME02_01 in response to Pfo(T3S) + *hopA1j* (Fig. S3), was matched with a strong transcriptional response probably due to HopA1j recognition (Fig. 5D). 1529 DEGs were upregulated in response to Pfo(T3S) + *hopA1j* treatment, and 1165 DEGs were downregulated. Consistent with moderate ion-leakage in *A. melanandra* upon Pfo(T3S) + EV and Psa3 Δ*33E* treatment (Fig. 1D, Fig. S3), 1105 DEGs were upregulated and 801 were downregulated in response to Pfo(T3S) + EV (Fig. 5E), and 389 DEGs are upregulated and 449 DEGs are downregulated in response to Psa3 V-13 Δ*33E* (Fig. 5C). When compared to the lack of a strong transcriptional response to Psa3 V-13 Δ*hrcC* in *A. melanandra*. This potentially suggests that *A. melanandra* may be recognising additional components beyond type III secreted effectors in Psa3 V-13 *33E* and Pfo(T3S) + EV that lack any effectors. These observations are consistent with callose deposition induced by Psa3 V-13, Psa3 Δ*33E*, Pfo(T3S) + *hopA1j* and Pfo(T3S) + EV on ME02_01 (Fig. 5F–G). Only the Psa3 V-13 Δ*hrcC* and the mock control did not induce callose deposition on ME02_01 (Fig. 5G), comparable to the lack of callose deposition in response to Psa3 V-13 treatment of the susceptible *A. chinensis* cultivar ‘Hort16A’ (Fig. 5G).

**Figure 5.**
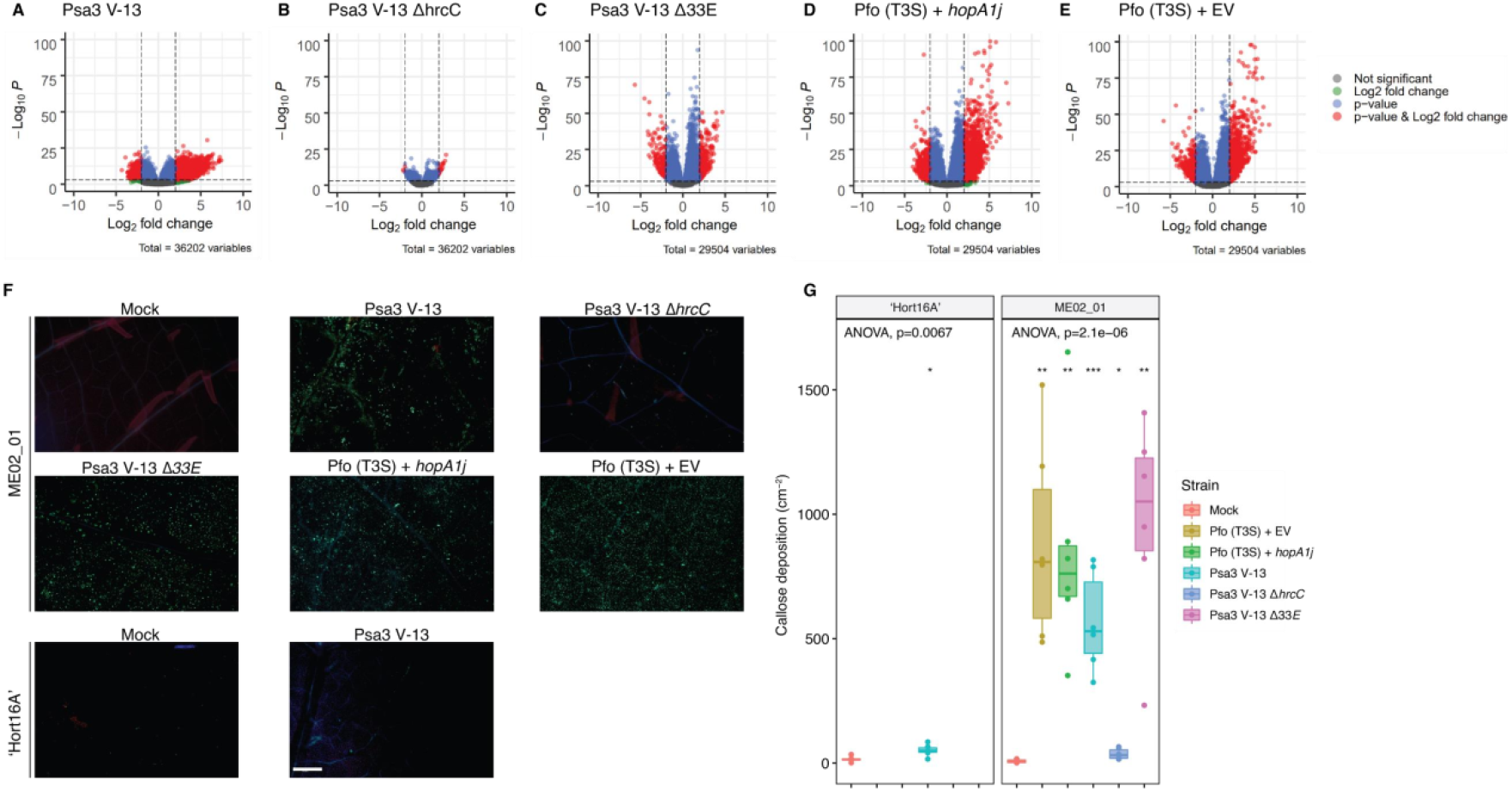
The magnitude of immune response of *Actinidia melanandra* ME02_01 to Pseudomonas *syringae* pv. *actinidiae* (Psa3 strains) or *P. fluorescens* (Pfo(T3S) strains). *A. melanandra* ME02_01 treated with (A) Psa3 V-13, (B) Psa3 V-13 *ΔhrcC*, (C) Psa3 V-13 Δ33E, (D) Pfo(T3S) + *hopA1i*, or (E) Pfo(T3S) + EV. Volcano plots of the differential expressed genes across different treatments (adjusted p-value <0.001, |log_2_ fold-change|>2). All data are from three biological replicates (n=3) with the 3, 6, 20-hour timepoints pooled. (F) Callose deposition induced by Pfo(T3S) carrying empty vector (EV) or the positive control *hopA1j*, wild-type Psa3 V-13, Psa3 V-13 Δ*hrcC* or Psa3 V-13 Δ*33E* in *A. melanandra* ME02_01 or *A. chinensis* var. *chinensis* ‘Hort16A’ leaves. The representative images were captured at 48⍰h after infiltration with mock (sterile ⍰11mM MgCl_2_) or bacterial strains. (G) The number of callose deposits per cm^2^ of leaf tissue from (F) was analysed with the ImageJ software. Box and whisker plots, with black bars representing the median values from six biological replicates and whiskers representing the 1.5 × inter-quartile range. Asterisks indicate significant differences from a one-way ANOVA and *post hoc* Welch’s t-test between the indicated strain and the mock treatment, where *p*≤0.05 (*), *p*≤0.01 (**), *p*≤0.001 (***), and *p*>0.05 (ns; not significant). All images were taken under similar conditions and magnification and the scale bar in the final panel represents 500 μm.

A heatmap visualisation of *A. melanandra* gene expression for the two early timepoints (3 and 6 hpi) indicates that Pfo(T3S) + *hopA1j*, Pfo(T3S) + EV and Psa3 V-13 Δ*33E*, share commonly up-regulated genes in cluster 1 represented by 3922 DEGs (Fig. 6A). The mid-timepoint (20 hpi) for Pfo(T3S) + *hopA1j* shares a similar response in this cluster with Psa3 V-13, indicating that their physiologically independent PTI + ETI responses converge, while PTI-only responses (Psa3 V-13 Δ*33E*, Psa3 V-13 Δ*hrcC*, and Pfo(T3S) + EV) are dampened as quickly and groups with mock-like response in cluster 2 with 4594 DEGs. Interestingly, the early response by ME02_01 to Psa3 V-13 and Psa3 V-13 Δ*hrcC*, appears to group together with little to no significant genetic response, indicating a lack of a strong PTI response by *A. melanandra*, supporting the lack of callose deposition seen in response to Psa3 V-13 Δ*hrcC* (Fig. 5G). Meanwhile, the strong callose deposition by Psa3 V-13 Δ*33E* and Pfo(T3S) + EV (Fig. 5G) is supported by a strong PTI-like/non-ETI type transient immune response not shown by Psa3 V-13 Δ*hrcC*, suggesting a recognition of the type III secretion system itself (Fig. 6A). Gene ontology term enrichment highlights the different responses to the Psa3 V-13, Psa3 V-13 Δ*33E* and Pfo (T3S) + EV treatments (Fig. S10). While protein phosphorylation and protein kinase activity were upregulated (particularly at 6 hpi) across all treatments, Psa3 V-13 Δ*33E* otherwise only weakly activated a response (Fig. S10). Nevertheless, for shared responses of defence-associated phosphorylation, phosphatase activity, vesicle transport and kinase activities, Psa3 V-13 Δ*33E* and Pfo (T3S) + EV shared early activation (3-6 hpi), with similar responses activated albeit slightly delayed at 6–20 hpi in Psa3 V-13 treatment. The strong non-host response of *A. melanandra* to both Psa3 V-13 and Pfo (T3S) + EV treatments was enriched for calcium ion binding, vesicle-mediated transport, salicylic acid mediated signalling, and programmed cell death. In contrast, photosynthesis and transmembrane transport appear to be downregulated in Psa3 V-13 Δ*33E* and Pfo (T3S) + EV, with the former and not the latter downregulated in response to Psa3 V-13 (Fig. S10).

**Figure 6.**
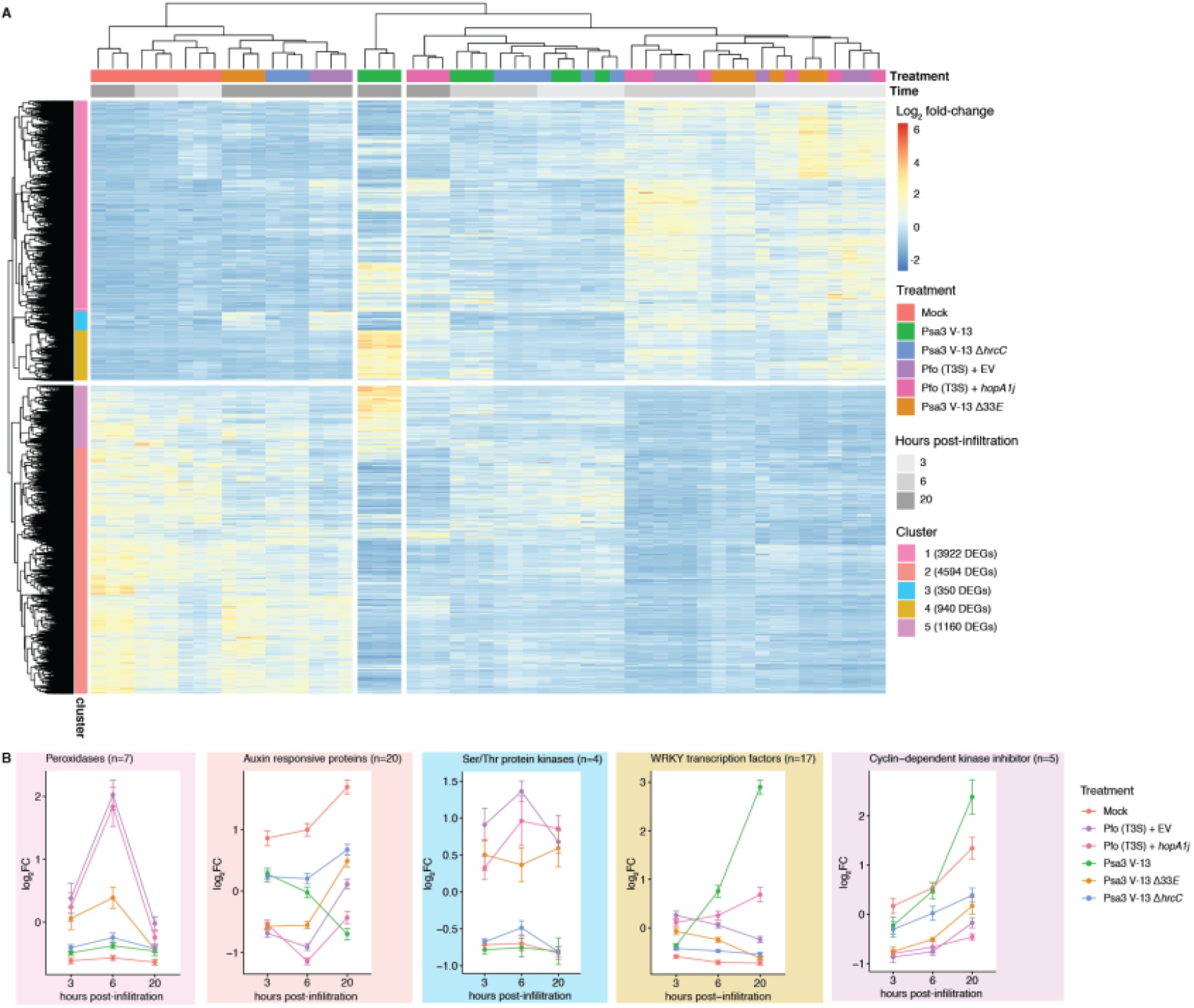
The genetic response of *Actinidia melanandra* ME02_01 is distinct between *Pseudomonas syringae* pv. *actinidiae* (Psa3 strains) and *P. fluorescens* (Pfo(T3S) strains). (A) Heatmap of differentially expressed genes induced by Psa3 V-13, Psa3 V-13 Δ*hrcC*, Psa3 V-13 Δ*33E*, Pfo(T3S) + *hopA1i*, and Pfo(T3S) + EV relative to mock (adjusted *p*-value <0.001). For each gene, raw counts were transformed by the median of ratios normalization and Z-score scaling. The pheatmap package was used to generate a heatmap, with genes divided into five clusters based on gene expression patterns using the hclust() function. (B) Expression of key differentially expressed genes by gene cluster in (A) for each treatment and timepoint. Error bars indicate standard errors.

Examining the clusters of gene expression in response to Psa3 V-13 strains in contrast to Pfo(T3S) strains, a clear segregation of PTI-like genes was seen, exemplified by peroxidases from cluster 1 that responded transiently to PTI-triggering both strains Pfo(T3S) and Psa V-13 Δ*33E* but not Psa3 V-13 and Psa3 V-13 Δ*hrcC* (Fig. 6B). Similarly, cluster 3 was represented by protein kinases that are probably activated during PTI (3-6 hpi) but repressed during ETI (Fig. 6B). Meanwhile, ETI-associated genes in cluster 4 responded only to the two ETI-triggering strains at 20 hpi, Psa3 V-13 and Pfo(T3S) + *hopA1j* (Fig. 6B). Cluster 2 represented the largest cluster of genes responsive to Psa3 V-13 treatment, whereby ETI appeared to suppress growth-associated genes including the Auxin responsive protein family, but surprisingly not affecting Pfo(T3S) + *hopA1j* and thus this cluster may represent a collection of genes affected by actions of Psa3 V-13 effectors (Fig. 6B). Taken together, these results suggest that the innate immune response to Psa3 V-13 (and Pfo(T3S) + *hopA1j*) is distinct from the somewhat transient immune response seen against other avirulent strains (Psa3 V-13 Δ*33E* and Pfo (T3S) + EV), none of which are is triggered by the T3S-lacking Psa3 V-13 Δ*hrcC*.

## Discussion

Psa3 strains that have lost some effectors have been isolated from several *Actinidia* species in the *Actinidia* species germplasm collections in Te Puke, New Zealand, where vines with diverse genotypes are co-located. In this setting, *A. melanandra* and *A. arguta* vines that recognise Psa3 can be located next to vines that do not recognise Psa3 and might not select for effector loss. Unsurprisingly, the emergence and spread of effector-loss strains are often limited to specific vines, where related strains can be isolated year after year. Interestingly, the effector-loss strains identified in this study were all obtained from a single ‘commercial’ block of *A. arguta* ‘HortGem’-series cultivars. In commercial *A. arguta* orchards, vines are always grown in monoculture and this may provide more opportunity for these strains to spread and ‘sweep’ the orchard population. Limited sampling of kiwiberry has been conducted to date (Vanneste et al., 2014). However, where an effector-loss strain has emerged in commercial orchards, these strains do not appear to spread across the whole orchard, nor between orchards (Hoyte et al., 2024).

Several studies have delved into the transcriptional response of *Actinidia* genotypes, varying in their susceptibility to Psa3, including immune response markers for ETI and PTI pathways (Qin et al., 2022; Song et al., 2019; T. Wang et al., 2018). In a similar vein, transcriptional reprogramming during Psa3-induced ETI in *A. melanandra* involved expression of canonical marker gene families (Fig. S7). These markers of PTI and ETI are well-documented in the scientific literature (Li et al., 2016; Peng et al., 2018; Yuan et al., 2021). This is consistent with cumulative evidence in model plants where PTI and ETI have many overlapping subsequent responses, differing predominantly in intensity and progression (Ngou et al., 2021; Yuan et al., 2021). The transcriptomic response to non-virulent Psa3 V-13 Δ*33E* and Pfo(T3S) + EV intensified between 3 and 6 hpi but waned from 20 hpi, suggesting that the PTI response in *A. melanandra* was transient, consistent with studies from model plants (Fig. 6; Bjornson et al., 2021; Li et al., 2016). The ETI response in *A. melanandra* was sustained in response to Psa3 V-13 (and mirrored by but delayed for Pfo(T3S) + *hopA1j*) (Fig. 6). Surprisingly, when compared with Psa3 V-13 and Pfo(T3S) + *hopA1j* (both expected and observed to trigger ETI) or Psa3 V-13 Δ*33E* and Pfo(T3S) + EV (both expected and observed to trigger PTI), Psa3 V-13 Δ*hrcC* stood out because of a lack of a significant genetic response and reduced intensity of expression (Figs 5 and 6). Similarly, callose deposition, a cell wall fortification response to pathogens, in challenged *A. melanandra* plants was prominent for all expected PTI and PTI + ETI treatments except in Psa3 V-13 Δ*hrcC* (Fig. 5F–G). *hrcC*– mutant strains lack a functional T3S assembly (Roine et al., 1997). This contrast between Psa3 V-13 Δ*hrcC* and Psa3 V-13 Δ*33E*/Pfo(T3S) + EV suggested that the T3S itself (or a component of it) is a salient feature in activating PTI in *A. melanandra*. Considering that recognition of Psa3 V-13 Δ*hrcC* response is also absent in ‘Hort16A’, this suggests a PTI-recognition of the Psa3 T3S by *Actinidia* spp. may be conserved (Jayaraman et al., 2021). From the kiwifruit breeding perspective, this study demonstrates the utility of Psa3 effector HopAW1 recognition as well as recognition of the Psa3 T3S for future mapping of resistance genes in the *A. melanandra* species against Psa3. To our knowledge, this is the first demonstrable opportunity to breed for gene-for-gene resistance in the Psa-kiwifruit pathosystem.

## Methodology

### Kiwifruit germplasm survey & Psa3 isolation

Samples were taken from leaf spots on vines in the *Actinidia* germplasm collections at the Plant & Food Research Te Puke and Kerikeri Research Orchards, as described in Hemara et al. (2022).

Quantitative PCR (qPCR) was carried out on an Illumina Eco Real-Time PCR platform (Illumina, Melbourne, Australia), following the protocol outlined by Andersen et al. (2017). Single colonies were tested with Psa-*ITS* F1/R2 PCR primers and primers specific to *hopZ5* to identify Psa3 strains. Samples that amplified in under 30 qPCR cycles were prepared as a 20% (w/v) glycerol stock for long-term storage.

### Psa3 DNA extraction & sequencing

Samples collected in 2017 and 2018 were extracted and sequenced as described in Hemara et al. (2022). For samples collected in 2022, DNA was purified using the Wizard® Genomic DNA Purification Kit (Promega). Libraries were constructed using the Seqwell Pureplex Unique Dual Index library preparation kit (SeqWell) and were sequenced on an Illumina MiSeq platform (paired-end 150 bp reads) (Illumina).

### Psa3 genome assembly, variant calling & effector presence analysis

Quality control reports for the raw sequencing reads were generated using FastQC. Snippy (version 4.6.0) was used to map reads to the reference genome of Psa3 V-13 (CP011972-3) and snippy-core was used to produce a core SNP alignment (Seemann, 2015). Gubbins (version 2.4.1) identified recombinant regions in this alignment, producing a filtered alignment of 391 bp (Croucher et al., 2015). RAxML (version 8.2.12; -f a -# 100 –m GTRCAT) was used to generate a maximum-likelihood phylogenetic tree with 100 bootstrap replicates (Stamatakis, 2014). The phylogeny and associated metadata were visualised with the R package ggtree (version 2.2.4; Yu et al., 2017). Only bootstrap support values of 50 or above were visualized.

Gene deletions were provisionally identified by CNVnator (version 0.4.1) using snippy’s .bam output of reads aligned to the Psa3 V-13 reference (Abyzov et al., 2011). Paired-end reads were assembled using shovill (version 0.9.0) (Seemann, 2019). Contigs were annotated with Prokka (version 1.3) (Seemann, 2014), preferentially using annotations from the Psa3 V-13 protein model. Pangenome analysis from Roary (version 3.7.0) (Page et al., 2015) identified effector gene presence and absence from the *de novo* assemblies.

### Data visualisation & statistical analysis

Statistical analysis was conducted in R (R Core Team, 2023), and figures were produced using the packages ggplot2 (Wickham, 2016) and ggpubr (Kassambara, 2017). Post-hoc statistical tests were conducted using the ggpubr (version 0.3.0) and agricolae (version 1.3) packages (de Mendiburu, 2017; Kassambara, 2017). The stats_compare_means() function from the ggpubr package was used to calculate omnibus one-way analysis of variance (ANOVA) or Kruskal-Wallis statistics to identify statistically significant differences across all treatment groups (Kassambara, 2017). For normally distributed populations, Welch’s t-test was used to conduct pair-wise parametric t-tests between an indicated group and a designated reference (Kassambara, 2017). For non-normal distributions, a Wilcoxon test was used to conduct pair-wise non-parametric tests between an indicated group and a designated reference (Kassambara, 2017). The HSD.test() function from the agricolae package was used to calculate Tukey’s Honest Significant Difference (de Mendiburu, 2017).

### Bacterial strains

Wild-type Psa3 isolates collected from Actinidia germplasm are described in Table S1. Psa3 V-13 effector knockout strains used in this study are described in Table S2. Psa3 V-13 Δ33E and Pfo(T3S) plasmid-complemented strains used in this study are described in Table S3.

All Psa3 and Pfo(T3S) strains were streaked from glycerol stocks onto LB agar supplemented with appropriate antibiotics; plates were sealed and grown for 48 h at 22°C. Overnight shaking cultures were grown in LB supplemented with appropriate antibiotics and incubated at 22°C with 200 rpm shaking. LB agar was supplemented with 12.5 μg/mL nitrofurantoin and 40 μg/mL cephalexin for Psa selection and 20 μg/mL chloramphenicol and 10 μg/mL tetracycline for Pfo(T3S) selection (all from Sigma Aldrich, Australia). To select for Psa3 and Pfo(T3S) strains carrying pBBR1MCS-5B vectors, LB agar was supplemented with 25 μg/mL gentamicin (Sigma Aldrich).

### Psa3 complete effector knockout

The Psa3 V-13 complete effector knockout strain was generated using the pK18mobsacB-based vectors used previously to generate single knockout strains (Hemara et al., 2022; Jayaraman et al., 2023). The single effectors or effector blocks were sequentially knocked out to generate the 33 effector knockout strain (Psa3 V-13 Δ*33E*) in the order: *hopZ5a/hopH1a (*using the *hopZ5a/hopH1a* double knockout vector*), hopBP1a (*previously *hopZ3), hopQ1a, hopAS1b, avrPto1b (*previously *avrPto5), avrRpm1a, fEEL (avrD1/avrB2b/hopF4a/hopAW1a/hopF1e/hopAF1b/hopD2a/hopF1a), hopF1c (*previously *hopF2), hopD1a (*using the *hopQ1a/hopD1a* double knockout vector*), CEL (hopN1a/hopM1f/avrE1d), hopR1b, hopAZ1a, hopS2b, hopY1b, hopAM1a-1, hopAM1a-2, hopBN1a, hopW1c (previously hopAE1), hopAU1a, hopI1c*, and *hopAH1* block *(hopAG1f/hopAH1c/hopAI1b*). Knockouts were confirmed by PCR and full genome sequencing of the Psa3 V-13 Δ*33E* strain to confirm all effectors were lost. Effectors that did not have a functional type III secretion signal owing to truncation or disruption, or did not possess a HrpL box promoter individually or in an operon (confirmed by expression analysis in (McAtee et al., 2018)) were not knocked out and included the following probable pseudogenic effector loci: *avrRpm1c, hopA1a, hopAA1b, hopAT1e (*previously *hopAV1), hopAB1b (*previously *hopAY1)*. The effectors *hopAA1d (CEL* block), and *hopAG1f/hopAI1b* (hopAH1c block) were also considered pseudogenes under these criteria, but were knocked out with other effectors in their block. Effector genes were plasmid-complemented back into Psa3 V-13 Δ*33E* following methodology established in Jayaraman et al. (2020).

### Callose deposition assays

Callose deposit quantification assays were conducted and images acquired as previously described in Jayaraman et al. (2023). Images were analysed using ImageJ software by determining the average area of a single callose deposit and then callose counts calculated based on total callose deposit area in each image.

### DAB staining

DAB assay was conducted as described in Jayaraman et al. (2021). Me02-01 leaves, vacuum infiltrated with Psa3 V-13, Psa3 V-13 Δ*hrcC* (resuspended in 10 mM MgCl2 and diluted to ∼10^8^ CFU/mL) and mock (10 mM MgCl2), were harvested and placed on a thin film of water in a sterile tub with lid covered for 48 h. Leaves were then vacuum infiltrated with DAB (3,3’-diaminobenzidine) solution (1 mg/mL DAB in 10 mM Na2HPO4 supplemented with 0.05% Tween® 20; all from Sigma Aldrich, Australia). Leaves were incubated in the dark for 6 h and de-stained with chloral hydrate solution (2.5 g per 1 mL water; Merck, NZ) for 24–48 h. Leaves were then washed with 70% ethanol and photographed.

### Pathogenicity assays

Pathogenicity assays were conducted as described in Hemara et al. (2022). Briefly, Actinidia tissue culture plantlets were flooded with Psa inoculum (10^6^ CFU/mL). Plantlets were grown in a climate control room at 20°C with a 16 h/8 h light/dark cycle. To quantify bacterial growth *in planta*, leaf disc samples were taken at 0 or 12 dpi.

### Ion leakage

Ion leakage was conducted as described in Hemara et al. (2022). Bacterial strains (Table S3) carrying empty vector or effector constructs were streaked from glycerol stocks onto LB agar plates with antibiotic selection, were grown for 2 days at 22°C, and were re-streaked onto fresh agar media, and allowed to grow overnight. Bacteria were then harvested from plates, were resuspended in 10 mM MgCl_2_, and were diluted to ∼10^8^ CFU/mL. Leaves were harvested from the tissue culture tubs and were submerged in 30 mL of bacterial inoculum. Vacuum-infiltrations were then carried out using a pump and glass bell. For each treatment, leaf discs (6 mm diameter) were harvested from the uniformly vacuum-infiltrated leaf area (6 cm) and were washed in distilled water for 1 h. Six discs were placed in 3 mL of water, and conductivity was measured over 48 h, using a LAQUAtwin EC-33 conductivity meter (Horiba). The standard errors of the means were calculated from five pseudobiological replicates.

### *Actinidia melanandra* DNA extraction & genome assembly

Leaves of *A. melanandra* (accession ME02_01) were collected from orchard-grown vines at the Plant & Food Research orchard, Te Puke, New Zealand. The nuclear genomic DNA was extracted with cetyltrimethylammonium bromide (CTAB)-based buffer as described previously (Naim et al., 2012). Hybrid whole genome sequence assembly of ME02_01 was developed using Oxford Nanopore Technologies PromethION data (5,599,455 base-called reads/45.3 Gb data with an estimated coverage of 59.64x) were assembled with Flye (version 2.9.3; Kolmogorov et al., 2019), 10x short-read data (283.00 million reads with an estimated coverage of 52.36x) were assembled with Supernova, 10x Genomics (Weisenfeld et al., 2017) and Hi-C (from Dovetail Genomics, Inc). Independent assemblies from Flye & Supernova were merged using Quickmerge (Chakraborty et al., 2016). Unassigned contigs with 20% or more alignment to existing mitochondrial or chloroplast genome sequences were separated to Fasta subsets containing 46 (mitochondria-related) and 15 (chloroplast-related) contigs. Assembly units were polished with NextPolish (Hu et al., 2020) and placed in linkages groups using aligned Hi-C reads (173.295 million reads mapped in pairs) and the YAHS assembler (version 1.2a.2). The YAHS assembly was critiqued using JuiceBox (Dudchenko et al., 2018). Contigs were placed in linkage groups. The chromosomes were subject to a single round of gap closure using abyss-sealer (Paulino et al., 2015) and Illumina paired-end reads, which closed 8.9% of the gaps. BRAKER3 (Gabriel et al., 2023) was used for gene predictions, along with RNAseq data (in the section below). Chimeric predictions (BRAKER3 prediction, which merged 2 or more separate genes into one prediction) were manually curated in WebApollo2 using physical evidence from RNA-Seq data as well as gene models from closely related *Actinidia* genomes aligned to the genome of ME01_01 using gmap (version 2023-04-28; Wu et al., 2016). The circos plot for the *A. arguta* and *A. melanandra* genomes was developed using Circos (version 0.23-1; Krzywinski et al., 2009) using the alignments performed by nucmer (using ‘—mum’) from MUMmer4 (version 4.0.0; Marçais et al., 2018) with alignment less than 10 kb filtered out.

### RNAseq and expression analyses

The RNAseq experiment was performed on tissue culture-grown plantlets of *A. melanandra* (accession ME02_01), kept at 22°C, 16 h/8 h light/dark cycle. Each tub contained three rooted plants spaced equally. Healthy plants aged between 24 and 36 days were used for the experiment with 8–12 leaves per plant from the top three internodes only. Complete plants were submerged into the respective treatments Psa3 V-13, Psa3 V-13 Δ33E, Psa3 V-13 ΔhrcC, Pfo(T3S) + hopA1j, or Pfo(T3S) + EV, resuspended in 10 mM MgCl_2_ at ∼10^8^ CFU/mL, or mock (10 mM MgCl_2_) and were vacuum infiltrated into the plants.

For each treatment, nine independent plants in three different tubs were harvested for each timepoint (3 h, 6 h, 10 h, 20 h, 30 h, and 40 h), with each biological replicate consisting of leaves from three separate plants. For Psa3 V-13 Δ33E, Pfo(T3S) + *hopA1j*, and Pfo(T3S) + EV, only three timepoints were harvested (3 h, 6 h, 20 h). Post-infiltration, only fully infiltrated leaves were harvested and snap-frozen in liquid nitrogen. Leaf tissues without infiltration (untreated) were also harvested just before the infiltration and labelled as 0 h. The full experimental procedure was performed three times, independently. RNA extraction was performed using the Spectrum™ Plant Total RNA Kit, quantified by spectrophotometry, and equal amounts of the total RNA were pooled up to 3 μg for mRNA short-read sequencing.

High-quality trimmed (≥Q30) PE (1,699.773 million clean reads/572 Gb data) and SE (582.08 million clean reads/58.7 Gb data) mRNA reads (100–150 bp length) were obtained for the samples (10–20 million reads per sample) using DNBSEQ™ G400 sequencing technology at BGI, Hong Kong, China and Illumina NovaSeq 6000 sequencing at Australian Genome Research Facility, Melbourne, Australia. The reads were assessed for quality using FastQC (Khetani, 2018) and aligned to the reference ME02_01 genome using HiSAT2 (Kim et al., 2019) and sorted BAMs were filtered for high-quality alignments. Feature counts were calculated from the alignments using the subread/1.5.3 package (Liao et al., 2014). For a few gene IDs, where more than one gene models were predicted, the gene ID row with the highest total counts was kept. If there was no significant difference between the alternate splice model, only one representative model was kept.

Differential expression analysis was performed between each bacterial treatment and the mock treatment. Differentially expressed genes (DEGs) were identified from gene counts using the Bioconductor package DESeq2 (version 1.28.0) (Love et al., 2014). Principal components analysis of normalised log_2_-transformed counts was performed using the estimateSizeFactors(), estimateDispersions() and varianceStabilizingTransformation() functions from DESEQ2 and the myPCA() function from the R package dataVisEasy. Log_2_-transformed DEGs with an adjusted p-value <0.001 were selected for further analysis. Enhanced volcano plots were produced using the lfcshrink() function from DESEQ2 and the R package EnhancedVolcano (version 1.6.0). Genes with an adjusted *p*-value <0.001 and |log_2_ fold-change|≥2 were considered as DEGs. For heatmap and cluster analysis, DEGs with an adjusted p-value of 0.001 were selected for further analysis irrespective of the magnitude of log_2_ fold-change. Heatmaps were generated with the R package pheatmap (version 1.0.12). Raw counts were normalised for heatmap visualisation using custom mor_normalization and cal_z_score functions (https://scienceparkstudygroup.github.io/rna-seq-lesson/06-differential-analysis/index.html). Gene expression clusters were created using the hclust() function. InterProScan (version 5.59-91.0) was used to provide gene ontology (GO) terms for ME02_01’s peptide sequences from the PANTHER, Pfam, and CDD databases (Jones et al., 2014). The AgriGO v2 Singular Enrichment Analysis (SEA) custom analysis tool was used to identify and compare GO term enrichment (Tian et al., 2017).

## Data Availability Statement

The data underlying this article are available in the GenBank Nucleotide Database (https://www.ncbi.nlm.nih.gov/genbank/) and Sequence Read Archive (https://www.ncbi.nlm.nih.gov/sra), including the Actinidia melanandra ME02_01 whole genome sequence (JBAMMV000000000) and experimental RNA sequencing reads (PRJNA1080659).

## Acknowledgements

The authors thank Sidney Scott for assistance with germplasm sampling, Juanita Dunn and Gavin Lloyd for assistance with research orchard access, and Dr Rebecca Bloomer and Dr Niels Nieuwenhuizen for reviewing the manuscript.

The authors acknowledge the use of New Zealand eScience Infrastructure (NeSI) high-performance computing facilities as part of this research. New Zealand’s national facilities are provided by NeSI and funded jointly by NeSI’s collaborator institutions and through the Ministry of Business, Innovation & Employment’s Research Infrastructure programme. URL https://www.nesi.org.nz.

