## Supplementary Material for "Identification and characterization of innate immunity in *Actinidia melanandra* in response to *Pseudomonas syringae* pv. *actinidiae*"

**Supplementary Figures**

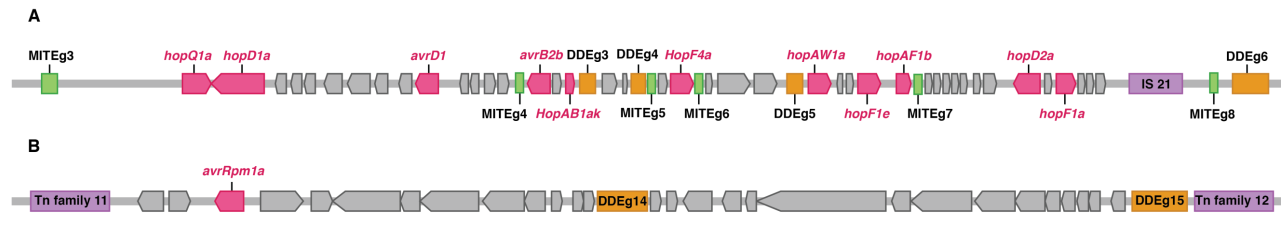

**Figure S1.** *Pseudomonas syringae* pv. *actinidiae* (Psa3) isolated from symptomatic germplasm *Actinidia* vines have gene deletions spanning recognized effectors. (A) The Psa3 X-469 gene deletion spans the effectors *hopQ1a, hopD1a, avrD1, avrB2b, hopAB1ak, hopF4a, hopAW1a, hopF1e, hopAF1b, hopD2a,* and *hopF1a*. (B) The Psa3 X-34 gene deletion spans the effector avrRpm1a and the conjugal transfer proteins on exapt ICE-A.

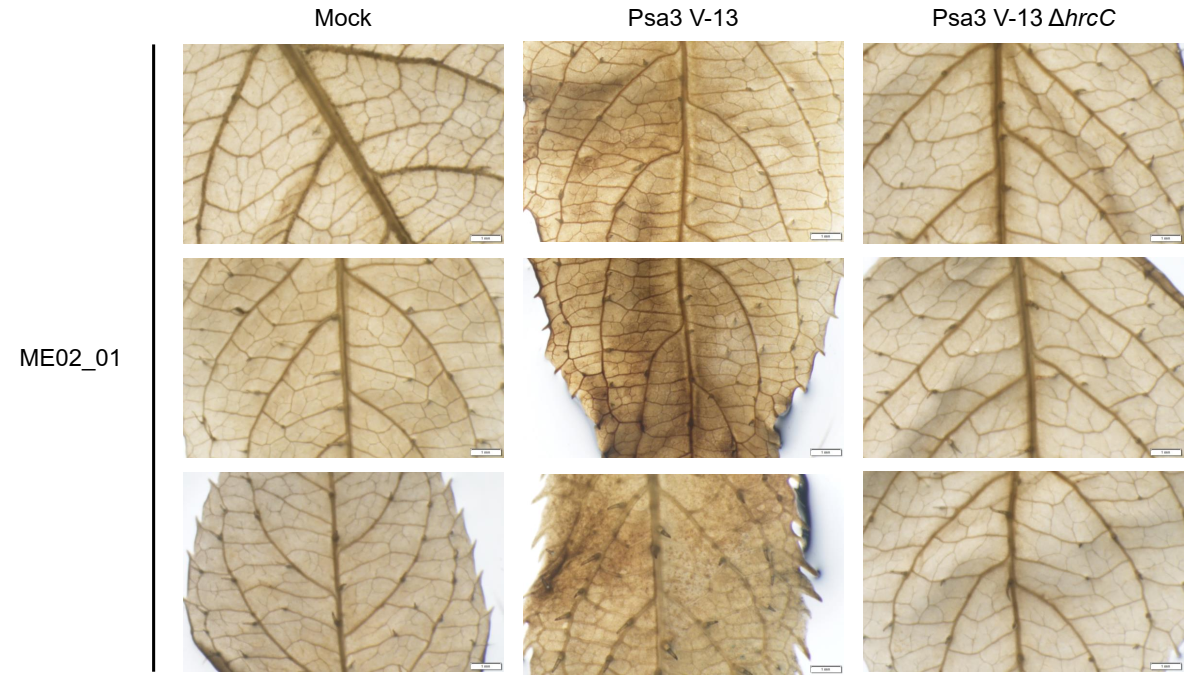

**Figure S2.** Reactive oxygen species (ROS) produced by *Actinidia melanandra* ME02_01 in response to *Pseudomonas syringae* pv. *actinidiae* (Psa3) V-13 but not Psa3 V-13 ∆*hrcC*. 3,3′-Diaminobenzidine (DAB) staining of ME02_01 leaves vacuum infiltrated with Psa3 V-13, Psa3 V-13 Δ*hrcC*, or buffer alone (mock) treatments (~10^8^ CFU/mL), DAB stained at 48 hpi and photographed at 10× magnification. Scale bars in each image represent 1 mm.

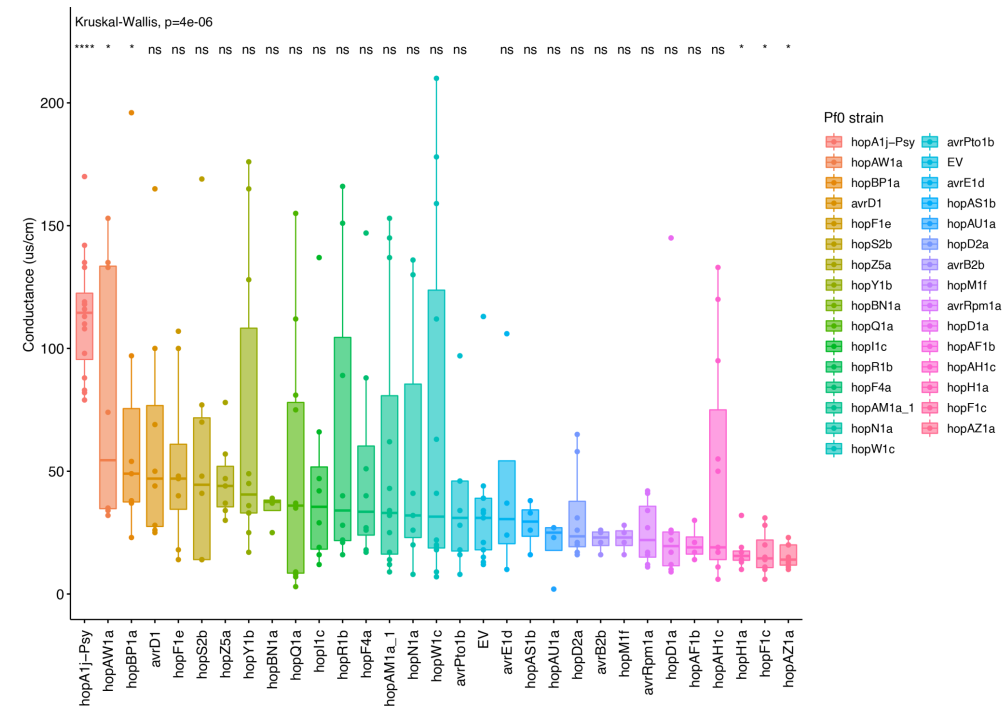
\

**Figure S3.** Multiple *Pseudomonas syringae* pv. *actinidiae* V-13 effectors trigger ion leakage in *Actinidia melanandra* ME02_01. *P. fluorescens* carrying a T3S from *P. syringae* pv. *syringae* 61 (Pfo(T3S)) was used to deliver individual Psa3 V-13 effectors into ME02_01 leaves by vacuum infiltration at ~10^8^ CFU/mL and ion leakage from cell death measured at 48 hours post infiltration. Asterisks indicate significant differences from a Kruskal-Wallis variance test and *post* *hoc* Wilcoxon rank-sum test between the indicated strain and Pfo(T3S) carrying an empty vector (EV), where *p*≤0.05 (*), *p*≤0.0001 (****), and *p*>0.05 (ns; not significant).

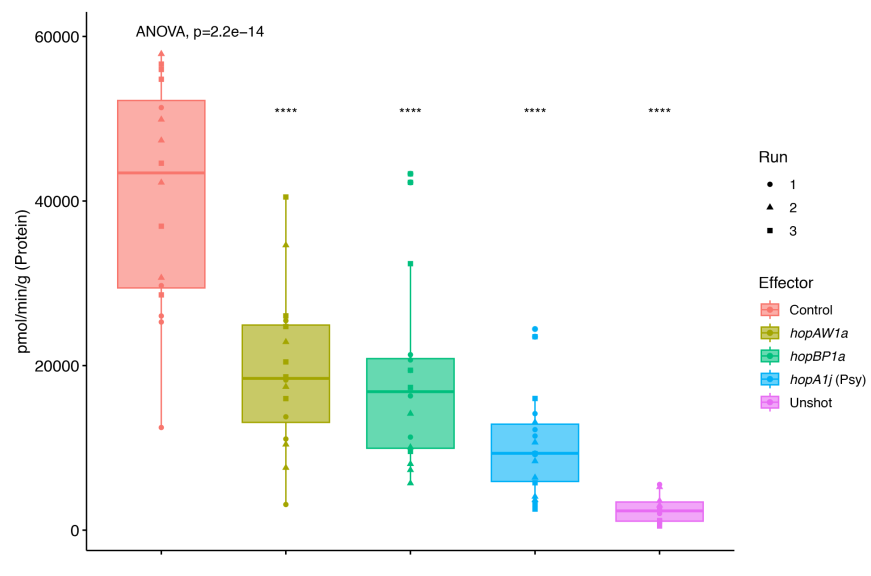

**Figure S4.** HopAW1a and HopB1a are recognized by *Actinidia melanandra* ME02_01 and trigger a reporter eclipse due to cell death. Effectors cloned in binary vector constructs tagged with GFP, or an empty vector (Control), were co-expressed with a β-glucuronidase (GUS) reporter construct using biolistic bombardment and priming in leaves from *A. melanandra* ME02_01 plantlets. The GUS activity was measured 48 hours after DNA bombardment. Error bars represent the standard errors of the means for two independent biological replicates with six technical replicates each (n = 12). HopA1j from *Pseudomonas syringae* pv. *syringae* 61 was used as positive control and un-infiltrated leaf tissue (Unshot) as a negative control. Asterisks indicate significant differences from a one-way ANOVA and *post hoc* Welch’s *t*-test between the indicated effector and no effector (control), where *p*≤0.0001 (****).

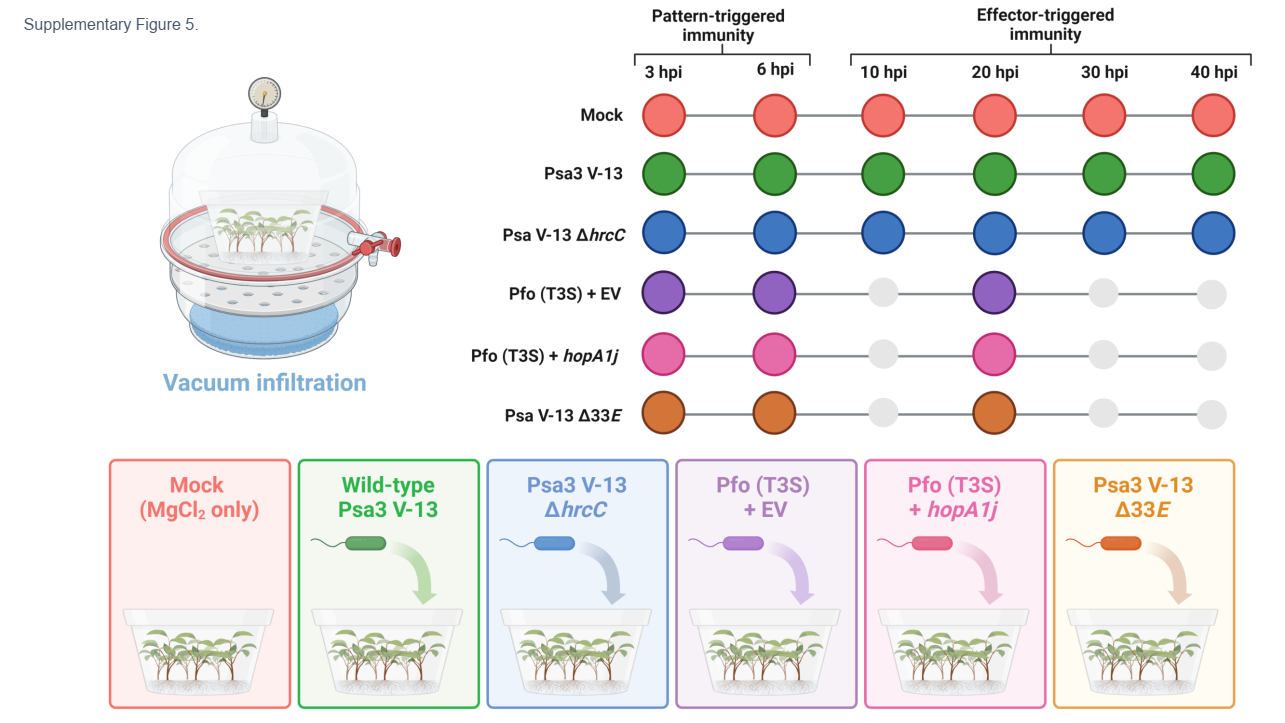

**Figure S5.** Experimental design. *Actinidia melanandra* ME02_01 was vacuum-infiltrated with mock (red), *Pseudomonas syringae* pv. *actinidiae* (Psa3) V-13 (green), Psa3 V-13 ∆*hrcC* (blue), Pfo (T3S) + EV (purple), Pfo (T3S) + *hopA1j* (pink), or Psa3 V-13 ∆*33E* (orange) at ~10^8^ CFU/mL and sampled over indicated key time periods, spanning pattern-triggered immunity (PTI) and effector triggered-immunity (ETI) windows.

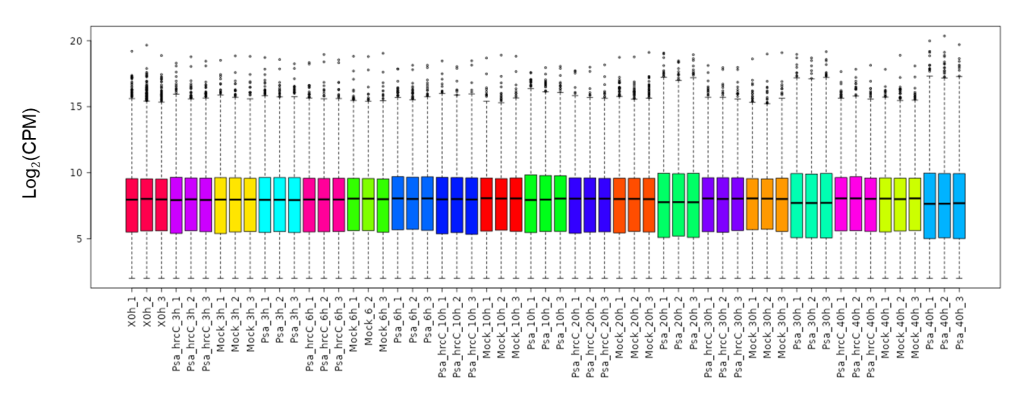

**Figure S6.** Normalized count data for RNAseq samples, Set 1. Log_2_CPM of samples for *Pseudomonas syringae* pv. *actinidiae* (Psa3) V-13, Psa3 V-13 Δ*hrcC*, and mock treatments harvested across all indicated 0–40 h timepoints.

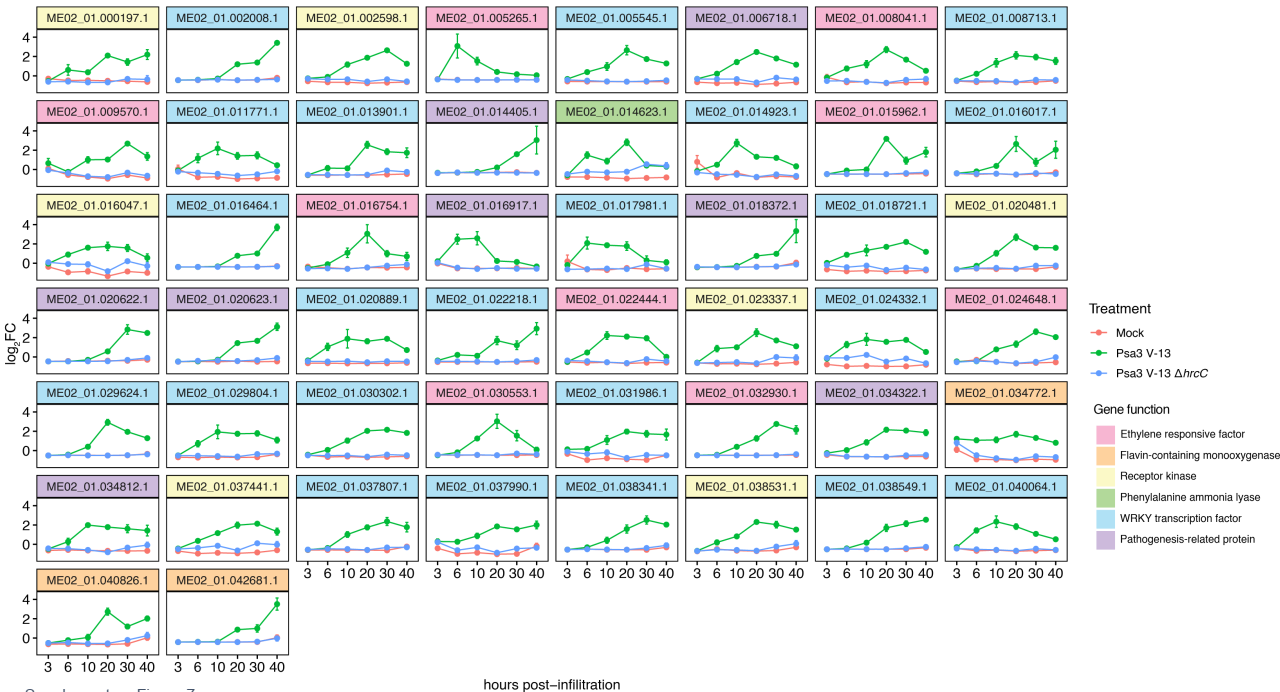

**Figure S7.** The transcriptional response of key marker genes to *Pseudomonas syringae* pv. *actinidiae* (Psa3) V-13, Psa3 V-13 ∆*hrcC*, or mock treatment in *Actinidia melanandra* ME02_01 over time. Log_2_ fold-change of key genes over the 40-h time series with each gene from a family of key marker genes indicated.

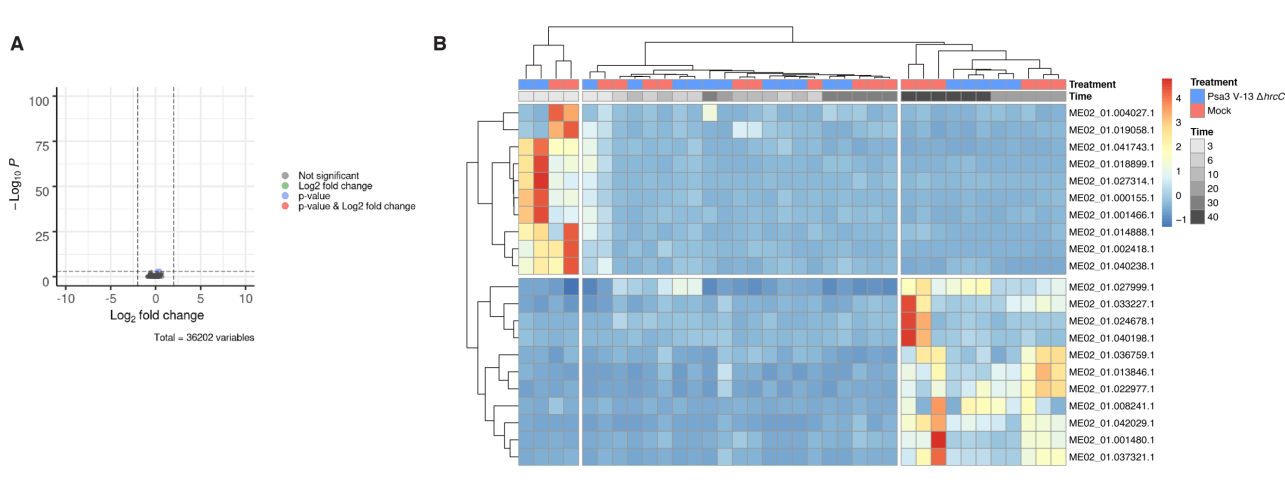

**Figure S8.** The transcriptional response to *Pseudomonas syringae* pv. *actinidiae* (Psa3) V-13 ∆*hrcC* treatment in *Actinidia melanandra* ME02_01 over time. **(A)** Volcano plots of the differential expressed genes (DEGs) from the Psa3 V-13 ∆*hrcC* on *A. melanandra* ME02_01, relative to mock (adjusted *p*-value <0.001, |log_2_ fold-change|>2). All data are based on three biological replicates (n=3) with the 3, 6, 10, 20, 30, and 40-hour timepoints pooled. **(B)** Heatmap of differential expression induced by Psa3 V-13 ∆*hrcC* treatment relative to mock (adjusted *p*-value <0.05).

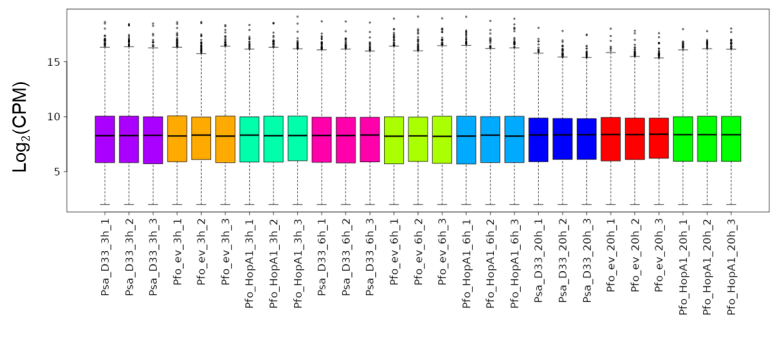

**Figure S9.** Normalized count data for RNAseq samples, Set 2. Log_2_CPM of samples for *Pseudomonas fluorescens* (Pfo(T3S)) + hopA1j, Pfo(T3S) + EV, and *P. syringae* pv. *actinidiae* (Psa3) V-13 ∆*33E* treatments harvested at 3, 6 and 20 h timepoints.

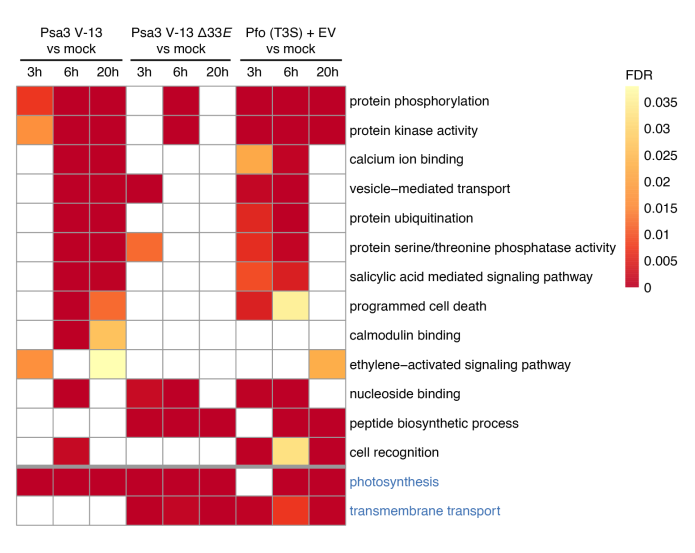

**Figure S10.** Gene ontology (GO) terms enriched in the differentially expressed subset of *Actinidia* genes for each treatment comparison and timepoint. The *p*-value represents the term enrichment at 3, 6, and 20 hpi. White cells are not enriched for a given GO term. Terms in blue text were downregulated, terms in black are upregulated.

**Supplementary tables**

**S1 Table. *Pseudomonas syringae* pv. *actinidiae* (Psa3) isolates from *Actinidia* germplasm.**

| Isolate | ICMP accession | Host | Year | Region |
| --- | --- | --- | --- | --- |
| Psa3 V-13 | ICMP 18884 | *Actinidia chinensis* var. *deliciosa* | 2010 | Bay of Plenty |
| Psa3 X_025 | ICMP 24617 | *Actinidia* spp. | 2017 | Bay of Plenty |
| Psa3 X_026 | ICMP 24618 | *Actinidia arguta* | 2017 | Bay of Plenty |
| Psa3 X_027 | ICMP 24619 | *Actinidia arguta* | 2017 | Bay of Plenty |
| Psa3 X_028 | ICMP 24620 | *Actinidia arguta* | 2017 | Bay of Plenty |
| Psa3 X_029 | ICMP 24621 | *Actinidia* spp. | 2017 | Bay of Plenty |
| Psa3 X_030 | ICMP 24622 | *Actinidia arguta* | 2017 | Bay of Plenty |
| Psa3 X_031 | ICMP 24623 | *Actinidia* spp. | 2017 | Bay of Plenty |
| Psa3 X_034 | ICMP 24626 | *Actinidia polygama* | 2017 | Bay of Plenty |
| Psa3 X_035 | ICMP 24627 | *Actinidia* spp. | 2017 | Bay of Plenty |
| Psa3 X_148 | ICMP 24628 | *Actinidia* spp. | 2018 | Bay of Plenty |
| Psa3 X_149 | ICMP 24629 | *Actinidia polygama* | 2018 | Bay of Plenty |
| Psa3 X_150 | ICMP 24630 | *Actinidia hypoleuca* | 2018 | Bay of Plenty |
| Psa3 X_151 | ICMP 24631 | *Actinidia arguta* | 2018 | Bay of Plenty |
| Psa3 X_152 | ICMP 24632 | *Actinidia arguta* | 2018 | Bay of Plenty |
| Psa3 X_153 | ICMP 24633 | *Actinidia polygama* | 2018 | Bay of Plenty |
| Psa3 X_154 | ICMP 24634 | *Actinidia arguta* | 2018 | Bay of Plenty |
| Psa3 X_155 | ICMP 24635 | *Actinidia polygama* | 2018 | Bay of Plenty |
| Psa3 X_156 | ICMP 24636 | *Actinidia chrysantha* | 2018 | Bay of Plenty |
| Psa3 X_157 | ICMP 24637 | *Actinidia arguta* | 2018 | Bay of Plenty |
| Psa3 X_158 | ICMP 24638 | *Actinidia arguta* | 2018 | Bay of Plenty |
| Psa3 X_159 | ICMP 24639 | *Actinidia eriantha* | 2018 | Bay of Plenty |
| Psa3 X_160 | ICMP 24640 | *Actinidia indochinensis* | 2018 | Bay of Plenty |
| Psa3 X_161 | ICMP 24641 | *Actinidia rufa* | 2018 | Bay of Plenty |
| Psa3 X_162 | ICMP 24642 | *Actinidia rufa* | 2018 | Bay of Plenty |
| Psa3 X_163 | ICMP 24643 | *Actinidia latifolia* | 2018 | Bay of Plenty |
| Psa3 X_165 | ICMP 24644 | *Actinidia arguta* | 2018 | Bay of Plenty |
| Psa3 X_166 | ICMP 24645 | *Actinidia arguta* | 2018 | Bay of Plenty |
| Psa3 X_167 | ICMP 24646 | *Actinidia arguta* | 2018 | Bay of Plenty |
| Psa3 X_168 | ICMP 24647 | *Actinidia arguta* | 2018 | Bay of Plenty |
| Psa3 X_169 | ICMP 24648 | *Actinidia arguta* | 2018 | Bay of Plenty |
| Psa3 X_170 | ICMP 24649 | *Actinidia arguta* | 2018 | Bay of Plenty |
| Psa3 X_171 | ICMP 24650 | *Actinidia arguta* | 2018 | Bay of Plenty |
| Psa3 X_172 | ICMP 24651 | *Actinidia arguta* | 2018 | Bay of Plenty |
| Psa3 X_461 |  | *Actinidia arguta* | 2022 | Bay of Plenty |
| Psa3 X_462 |  | *Actinidia arguta* | 2022 | Bay of Plenty |
| Psa3 X_463 |  | *Actinidia arguta* | 2022 | Bay of Plenty |
| Psa3 X_464 |  | *Actinidia arguta* | 2022 | Bay of Plenty |
| Psa3 X_465 |  | *Actinidia arguta* | 2022 | Bay of Plenty |
| Psa3 X_466 |  | *Actinidia arguta* | 2022 | Bay of Plenty |
| Psa3 X_467 |  | *Actinidia arguta* | 2022 | Bay of Plenty |
| Psa3 X_468 |  | *Actinidia arguta* | 2022 | Bay of Plenty |
| Psa3 X_469 | ICMP 25124 | *Actinidia arguta* | 2022 | Bay of Plenty |
| Psa3 X_470 |  | *Actinidia arguta* | 2022 | Bay of Plenty |
| Psa3 X_471 |  | *Actinidia arguta* | 2022 | Bay of Plenty |
| Psa3 X_472 |  | *Actinidia arguta* | 2022 | Bay of Plenty |
| Psa3 X_473 |  | *Actinidia arguta* | 2022 | Bay of Plenty |
| Psa3 X_474 |  | *Actinidia arguta* | 2022 | Bay of Plenty |
| Psa3 X_475 | ICMP 25125 | *Actinidia hypoleuca* | 2022 | Bay of Plenty |
| Psa3 X_476 |  | *Actinidia arguta* | 2022 | Bay of Plenty |
| Psa3 X_477 | ICMP 25126 | *Actinidia melanandra* | 2022 | Bay of Plenty |
| Psa3 X_478 |  | *Actinidia polygama* | 2022 | Bay of Plenty |
| Psa3 X_479 | ICMP 25127 | *Actinidia glaucophylla* | 2022 | Bay of Plenty |
| Psa3 X_480 |  | *Actinidia arguta* | 2022 | Kerikeri |
| Psa3 X_481 |  | *Actinidia latifolia* | 2022 | Kerikeri |
| Psa3 X_483 |  | *Actinidia arguta* | 2022 | Kerikeri |
| Psa3 X_484 |  | *Actinidia callosa* var. *henryi* | 2022 | Kerikeri |
| Psa3 X_485 |  | *Actinidia arguta* | 2022 | Kerikeri |

**S2 Table. *Pseudomonas syringae* pv. *actinidiae* (Psa3) V-13 knockout strains used in this study.**

| **Strain** | **Description** | **Source** |
| --- | --- | --- |
| Psa3 V-13 Δ33*E* | deleted 33 effectors | This study |
| Psa3 V-13 Δ*hrcC* | deleted *hrcC* | (Straub et al., 2018) |
| Psa3 V-13 Δ*hopAW1a* | deleted *hopAW1a* | (Hemara et al., 2022) |
| Psa3 V-13 Δ*hopZ5a* | deleted *hopZ5a* | (Hemara et al., 2022) |
| Psa3 V-13 Δ*avrRpm1a* | deleted *avrRpm1a* | (Hemara et al., 2022) |
| Psa3 V-13 Δ*hopF1c* | deleted *hopF1c* and associated *shcF* | (Hemara et al., 2022) |

**S3 Table. *Pseudomonas syringae* pv. *actinidiae* (Psa3) V-13 ∆33*E* and *P. fluorescens* plasmid-complemented strains used in this study.**

| **Strain** | **Description** | **Source** |
| --- | --- | --- |
| Psa3 V-13 Δ33*E* + EV | Plasmid-complemented with empty vector (pBBR1MCS-5) | This study |
| Psa3 V-13 Δ33*E* + pBBR1MCS-5:*hopAW1a* | Plasmid-complemented with *hopAW1a* (cloned under native promoter) | This study |
| Psa3 V-13 Δ33*E* + pBBR1MCS-5:*hopZ5a* | Plasmid-complemented with *hopZ5a* (cloned under native promoter) | This study |
| Psa3 V-13 Δ33*E* + pBBR1MCS-5:*avrRpm1a* | Plasmid-complemented with *avrRpm1a* (cloned under native promoter) | This study |
| Psa3 V-13 Δ33*E* + pBBR1MCS-5:*hopF1c* | Plasmid-complemented with *hopF1c* (cloned under native promoter) | This study |
| Pf0 (T3S) + EV | Plasmid-complemented with empty vector (pBBR1MCS-5) | (Jayaraman et al., 2020) |
| Pf0 (T3S) + pBBR1MCS-5:*hopA1j* | Plasmid-complemented with *shcA* and *hopA1j* from *P. syringae* pv. *syringae* 61 | (Jayaraman et al., 2021) |
